# Pro-tumoral Ca^2+^ signaling is dependent on Ca-α1T and Slowpoke channels in *Drosophila melanogaster* glioma

**DOI:** 10.1101/2025.07.16.665071

**Authors:** Lía Alza, Patricia Montes-Labrador, Diego Megías, Andreu Casali, Sergio Casas-Tintó, Judit Herreros, Carles Cantí

## Abstract

**Background:** Ion channel-mediated cytosolic Ca^2+^ oscillations play a crucial role in promoting the growth of glioblastoma, a tumor of poor prognosis. Here, we have studied the roles of Ca-α1T and Slowpoke ion channels in a *Drosophila melanogaster* glioblastoma model. Although their mammalian orthologs have been implicated in glioblastoma cell viability *in vitro*, their oncogenic potential remains uncharacterized in a complex *in vivo* context.

**Results:** We show that glial-specific RNAi targeting *Ca-α1T* and *Slowpoke* in gliomas reduces glioma cell number and membrane extension and decreases glioma cell Ca²⁺ activity and downstream activation of PI3K and pRIII/pERK signaling. Moreover, *Slowpoke* knockdown significantly extends glioma fly survival to near-control values and improves glioma-associated neurodegeneration. RNAseq transcriptomic analysis reveals shared regulation by both channels of pathways involved in metabolic rewiring, cell adhesion and excitatory neurotransmission, while uncovering distinct effects on synaptic function.

**Conclusions:** Both Ca-α1T and Slowpoke are essential for pro-tumorigenic Ca²⁺-dependent signaling in *Drosophila* gliomas. Common effects observed upon gene knockdown support a coordinated function for these channels at the plasma membrane. Nevertheless, their distinct impacts on survival and gene expression profiles highlight non-redundant roles. Notably, *Slowpoke* knockdown promotes increased survival and neuroprotection, associated with the repression of synaptic genes aberrantly upregulated in gliomas, thereby identifying Slowpoke as a promising therapeutic target in glioblastoma.

## Background

Gliomas are primary brain tumors classified by the World Health Organization (WHO) from grade I to IV according to their malignancy. Glioblastoma (GBM) is the most frequent and aggressive type of glioma, which, despite surgery, chemotherapy and radiotherapy remains an incurable disease [1]. The understanding of this tumor is rapidly evolving with the recognition of the involvement of neural elements in its progression [2, 3]. As novel molecular targets are emerging against the disease, there is the need for using preclinical model systems to study their functions in a complex context [4].

Ca^2+^ is a universal second messenger involved in transcriptional gene regulation and activation of different cell processes determinant for cell fate and viability. Cancer cells consistently experience a remodeling of Ca^2+^-dependent pathways promoting cell proliferation and survival, including the aberrant expression of ion channels which enable Ca^2+^ influx from the extracellular milieu [5, 6]. Among them, voltage-gated Ca^2+^ channels from the CaV3 family have been proposed as prognostic markers and therapeutic targets against GBM [7–9]. Remarkably, different studies have shown that CaV3 gene silencing or application of pharmacological blockers induces either GBM cell cycle arrest or apoptosis, although the effects of the latter cast doubt over their specificity [10–13].

While the contribution of CaV3 channels to GBM cell viability *in vitro* is well documented, their roles in preclinical models have only been partially addressed [14]. Targeting CaV3 in the tumoral setting may have effects on the tumor environment that have not been addressed *in vitro.* Furthermore, the *modus operandi* of CaV3 channels in the steadily depolarized plasma membranes of cancer cells remains unclear. In this respect, we have hypothesized that CaV3 channels act in concert with Ca^2+^-activated K^+^ channels (KCa channels) to promote membrane hyperpolarization, thus favoring Ca^2+^ influx through multiple Ca^2+^ channel types and enabling G1 to S cell cycle progression [15]. As a matter of fact, diverse studies have shown that KCa channels crucially regulate GBM cell growth and migration [16–18]. Recently, it has also been reported that the KCa1.1 isoform plays a role in cell viability of BRAF-mutant GBM cells and drives proliferation in a paediatric glioma model [19]. Interestingly, a splice variant of KCa1.1 displaying an enhanced Ca^2+^ sensitivity (namely the gBK isoform) is overexpressed in gliomas compared to non-malignant human cortical tissues, and tumor malignancy positively correlates with gBK expression [20]. Yet, the relevance of KCa channels is not limited to promoting autonomous GBM cell proliferation, survival and invasiveness. The present view of GBM is that of a functional adaptable cell network, including tumor cells that actively communicate with other tumoral cells, glia and neurons surrounding the tumor [21–23]. Network connections are established by long membrane protrusions, known as tumor microtubes (TM) [24]. In this context, KCa3.1 is known to regulate the propagation of intracellular Ca^2+^-waves along TM and to promote GBM growth through MAPK and NFkB pathways [25, 26].

In this study, we set on investigating the functional roles of Ca-α1T (CaV3 ortholog) and Slowpoke (Slo, KCa1.1 ortholog) in a *Drosophila melanogaster* glioma model to address their contribution to GBM growth and progression *in vivo*. Previous studies have reported *Ca-*α*1T* expression in neurons across various regions of the adult fly brain, where it plays a pro- awakening role (Jeong et al., 2015). Regarding *Slo*, it is broadly expressed in the *Drosophila* nervous system, in which it undergoes tissue and developmental-specific splicing to regulate neuronal firing pattern and neurotransmitter release [27–30]. The variety of genetic tools available and the high degree of homology between the fly and the human genomes makes *Drosophila* glioma an appealing model, where new findings on the GBM pathobiology have been generated or validated [22, 31, 32]. Fly gliomas are induced by the overexpression of constitutively active Epidermal Growth Factor Receptor (*EGFR*) and Phosphoinositide 3-kinase (*PI3K*) in glial cells [33], mirroring the most prevalent mutations found in GBM patients [34]. Our results from RNAi-expressing glioma flies indicate that both ion channels engage in glioma cell proliferation by augmenting Ca^2+^ activity and participating in PI3K/AKT and ERK pathways.

Notably, we find that glioma-induced neuronal degeneration and fly survival improve by reducing *Slo* expression in gliomas, but not *Ca-*α*1T* that has noxious effects in control flies. In addition, RNAseq transcriptomics indicate that both channels are involved in the metabolic reprogramming of glioma cells and the promotion of glutamatergic signaling, substantiating their pivotal role in the glioma neuroscience landscape.

## Results

### Ca-α1T and Slo are involved in *Drosophila* glioma cell proliferation and glial membrane extension

In our *Drosophila* glioma model, glioma-forming mutations (constitutively active forms of *EGFR^λ^*and *dp110^CAAX^*) are transcribed in glial cells simultaneously with the interfering RNAs (RNAi) targeting either the calcium channel *Ca-α1T* or the KCa potassium channel *Slo* encoding genes. This strategy was designed to explore the contribution of these ion channels to glioma cell proliferation while the oncogenic drivers remain active. Moreover, this model allows the visualization of glial cell membranes as myristoylated *RFP (mRFP)* is expressed under the glial-specific *Repo* promoter.

We used the Minos Mediated Integration Cassette (MiMIC) system, which utilizes a transposon encoding a GFP that integrates downstream of the target gene promoter [35] [40]to quantify the expression of Ca-α1T and Slo GFP fusion proteins in both control and glioma brains. We developed a customized ImageJ macro to quantify GFP intensity on mRFP-glial membranes (Table S1). We observed that *Ca-α1T* is expressed in tumoral and control brains, with a slightly higher expression in glioma cells (Fig. S1A and B). In contrast, *Slo* is expressed in control and tumoral brains at similar levels (Fig. S1C and D).

We next validated the RNAi efficiency of the constructs chosen to target *Ca-α1T* and *Slo* in glia by performing qPCR of whole larvae expressing either *Ca-α1T*-RNAis or *Slo*-RNAis under the control of the *tubulin* promoter. The efficiency of the gene silencing by all RNAi strains used was confirmed (Fig. S2). Furthermore, knockdown of *Slo* was also confirmed by immunocytochemistry using an anti-Slo antibody in *Slo* RNAi-expressing gliomas (Fig. S2C).

Having confirmed gene knockdown by the RNAis, we analyzed glial cell proliferation in control gliomas vs. gliomas expressing *Ca-α1T* or *Slo-RNAi*. Brain lobes of third-instar larvae were immunostained with an anti-Repo antibody. Gliomas exhibited a significantly higher number of glial cells per lobe compared to control brains, as expected due to activation of EGFR and PI3K oncogenic cascades (Fig. 1). Two different RNAi constructs were employed to target. In contrast, expression of *Ca-α1T* or *Slo-*RNAis in gliomas reduced glial cell counts to values similar to control brains (Fig. 1 and S3). Remarkably, knocking down either channel in control brains did not change the number of glial cells compared to controls, indicating that their cell viability-promoting effects were specific to tumoral glia (Fig. 1B and C), with the only exception of the *Slo-*RNAi(B) line that significantly reduced control glial numbers (Fig. S3C). Interestingly, the reduction in glial cell numbers correlated with the decrease of the diameter of the glioma brain lobes upon *Ca-α1T* or *Slo-*RNAi expression (Fig. S3D and E).

**Fig. 1.**
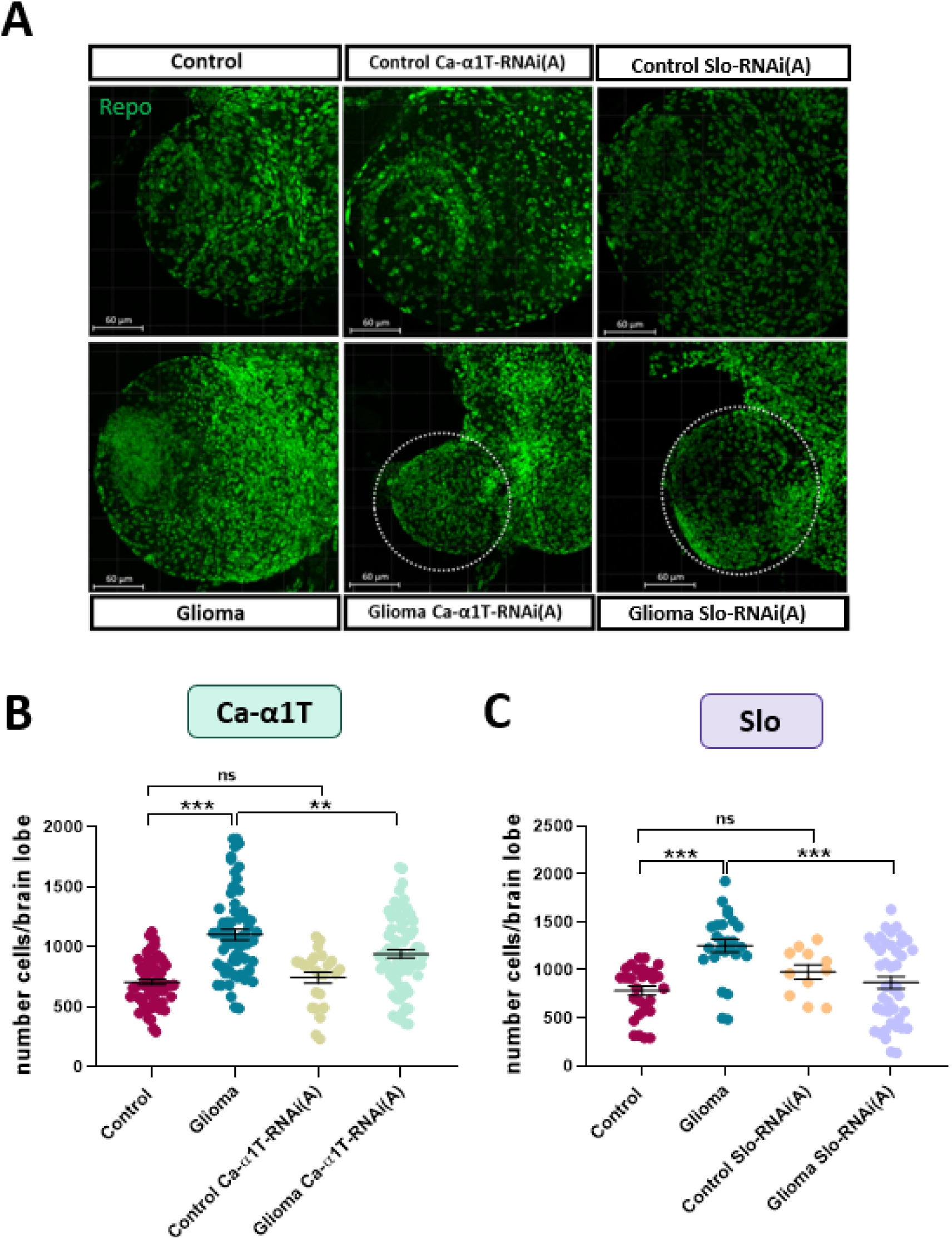
Knocking down Ca-α1T and Slo reduces glioma cell proliferation **(A)** Representative confocal images of Drosophila larval brain lobes showing glial cells immunostained for Repo. Scale bar: 60 μm. **(B-C)** Quantification of glial cell numbers. Glioma significantly increased glial cell counts compared to control. Reducing Ca-α1T (B) or Slo (C) levels in gliomas significantly decreased glial cell numbers compared to control gliomas, but did not alter the number of glial cells in control brains.

Glial membrane extension, indicative of TM expansion, is part of the neurodegenerative process that glioma cells impinge on the brain and correlates with worse prognosis in human patients. We leveraged that our model expresses mRFP in cell membranes, to analyze glial membrane extension in gliomas and gliomas expressing *Ca-α1T* or *Slo*-RNAis. mRFP quantification showed a significant increase in glial membrane volume of glioma brain lobes compared to controls (Fig. 2), consistent with prior studies (Portela et al., 2019a). In contrast, knocking down *Ca-α1T* or *Slo* in glioma brains significantly reduced glial membrane volume, restoring it to levels comparable to those in control brains (Fig. 2B and C). Importantly, we also observed a reduction in the membrane volume per cell ratio, indicating that the decrease in membrane extension was not solely due to the reduction in glial cell number (Fig. 2D and E). Notably, we did not observe differences in membrane extension upon *Ca-α1T* or *Slo-*RNAi *expression* in control brains were observed (Fig. 2B and D), suggesting that both ion channels are specifically involved in the extension of tumoral membranes. Glioma membrane results were reproduced by a second RNAi for either channel (Fig. S4).

**Fig. 2.**
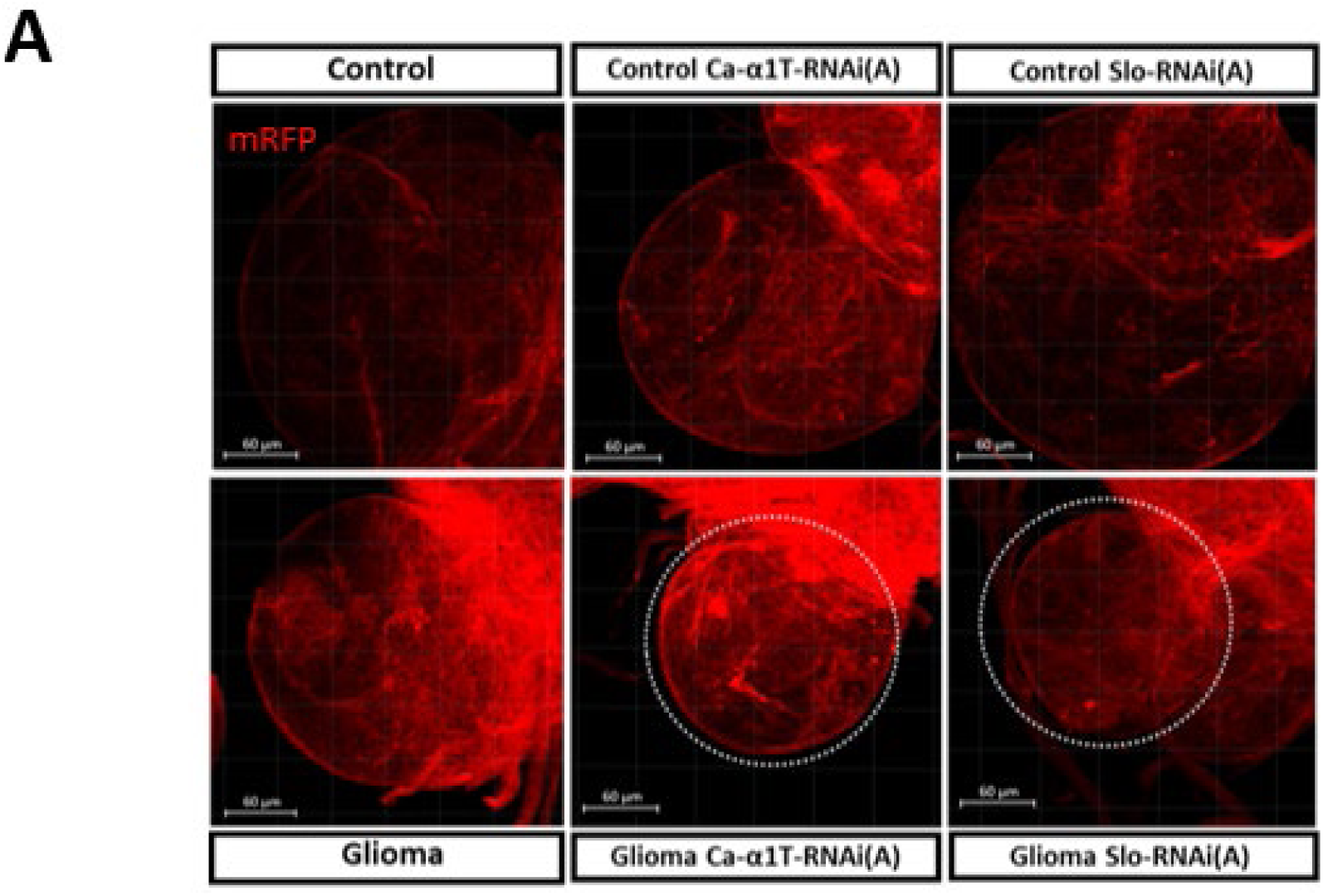
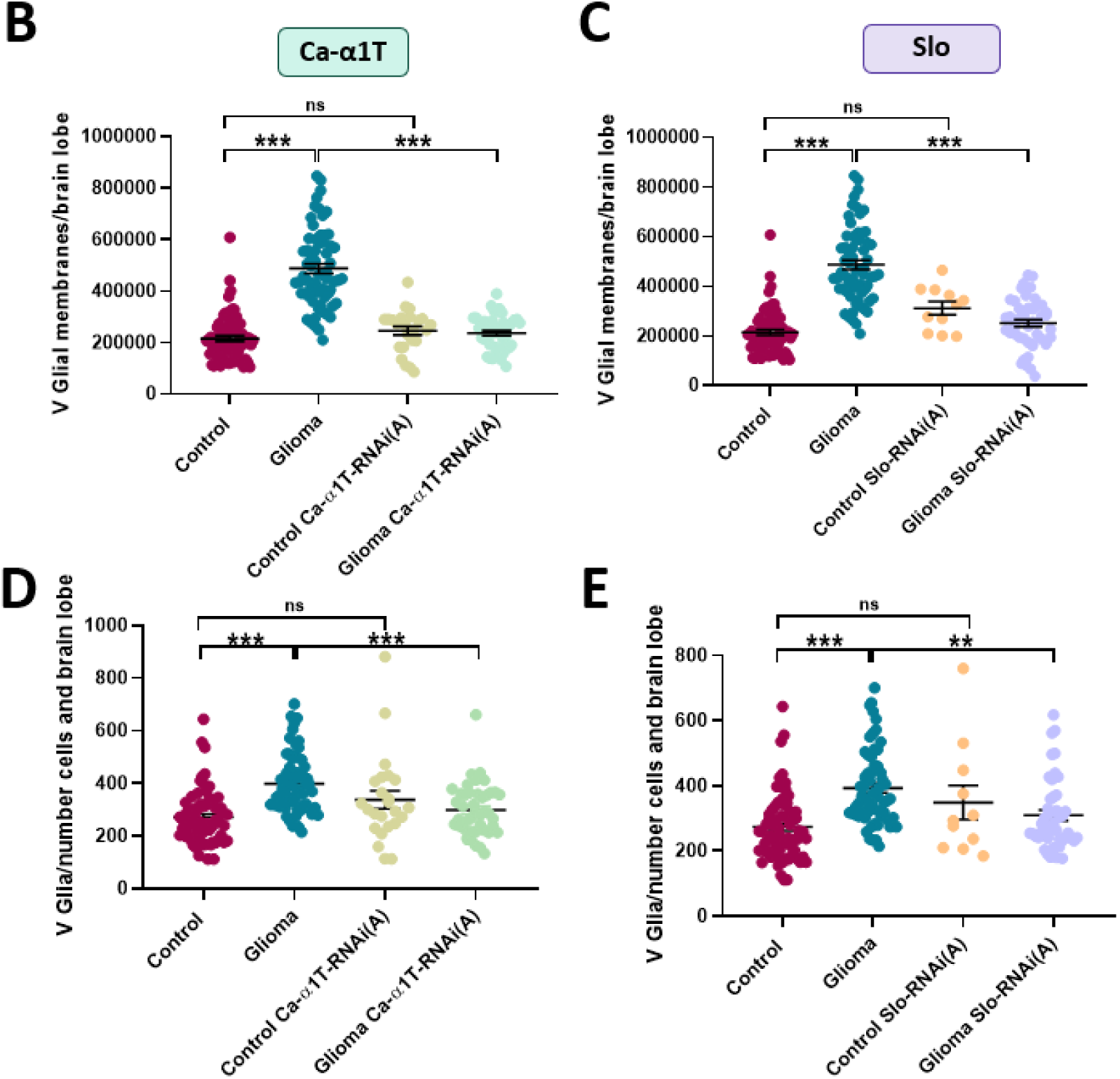
Knocking down Ca-α1T or Slo reduces glial membrane extension in Drosophila glioma cells. **(A)** Representative confocal images of glial membranes in red (mRFP) in Drosophila larval brain lobes. Scale bar: 60 µm. (**B-C)** Quantification of glial membrane volume per cell. Gliomas exhibited significantly greater glial membrane volume per cell unit than control brains. Knocking down Ca-α1T or Slo significantly reduced membrane volume per cell in gliomas but not in control brains. **(D-E)** Quantification of glial membrane volume per brain lobe. Gliomas showed an increased glial membrane volume compared to controls. Knocking down Ca-α1T or Slo decreased membrane volume of glioma cells but not of non-tumoral glia. Values represent means and SEM derived from more than three independent experiments, with a sample size between n=11-76 brain lobes. Multiple comparisons with Tukey’s test (**p<0.01, ***p<0.001).

### Ca-α1T and Slo participate in pRII- and PI3K-activated pathways

Having observed the reduction in glioma cell number in gliomas expressing *Ca-α1T* or *Slo-* RNAis, we reasoned that these channels might participate in the hyperactivated pathways driving glioma progression: EGFR and PI3K. We first assessed the levels of phosphorylated Rolled, pRll (the phosphorylated form of the ortholog of human ERK1/2) that results from the activation of EGFR/RAS pathway. As previously described [33], pRII immunostaining increased in glioma brains compared to control brains (Fig. 3A and B). However, *Ca-α1T* or *Slo-*RNAi expression in gliomas significantly reduced pRll immunostaining (Fig. 3A and B), suggesting that these channels are involved in RII activation in *Drosophila* gliomas.

**Fig. 3.**
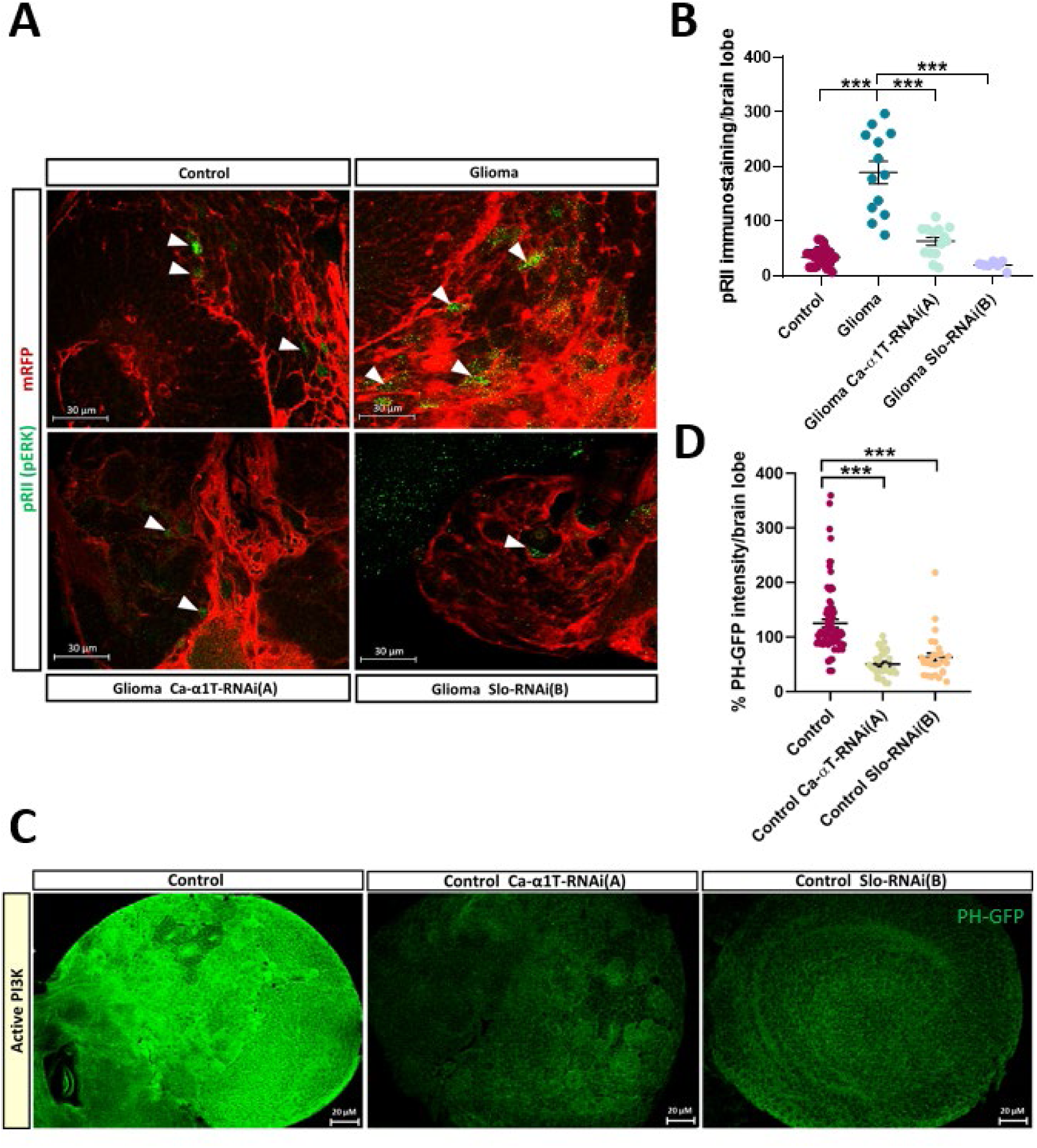
Ca-α1T and Slo knockdown reduce pRll and PI3K activation. **(A)** Representative confocal images showing pRII (pERK) immunostaining (green) in glial cells (glial membranes labelled in red with mRFP). Scale bar: 20 µm. **(B)** Quantification of pRll-positive glial cells (surrounded by mRFP). Glioma significantly increased pRll compared to control brains. Knocking down Ca-α1T or Slo in glioma reduced pRll-positive cells compared to control gliomas. One-way ANOVA with Tukey’s multiple comparisons test (*p<0.05, ***p<0.001). **(C)** Representative images of GFP fluorescence corresponding to PI3K activation revealed by the PH-GFP construct in control brains or brains knocked down for each ion channel. Scale bars: 20µm. **(D)** Quantification of GFP intensity per brain lobe compared to control brain levels. Reduced Ca-α1T or Slo expression in control brains significantly decreased PI3K activation. T-Test (***p<0.001, *p<0.05). Data represents mean and SEM from at least three independent experiments and a sample size of n=7-25 brain lobes per condition for pRll and n=21–35 brain lobes for PH-GFP.

To analyze the possible role of *Ca-α1T* and/or *Slo* in the PI3K pathway, we used a Pleckstrin Homology domain-Green Fluorescent Protein (PH-GFP) as a reporter of PI3K activity [37]. Since this reporter is ubiquitously expressed, quantification was conducted across the entire brain. These experiments were performed using control flies in which *PI3K* expression was not forced. Knocking down *Ca-α1T* or *Slo* significantly decreased PI3K activation in the brain (Fig. 3C and D), indicating that both channels take part in the PI3K pathway.

### Ca-α1T and Slo support heightened Ca^2+^ activity in glioma

Glioma network relies on synchronized Ca^2+^ flux through TM that drive tumor growth [26]. To understand the contribution of *Ca-α1T* and *Slo* in glioma pathophysiology, we employed the Ca^2+^-dependent nuclear import of LexA (CaLexA) reporter system, a robust tracer of neural activity [36]. CaLexA GFP fluorescence quantification revealed a 2.24-fold increase of Ca^2+^ activity in gliomas compared to control brains (Fig. 4). Interestingly, knockdown of either *Ca- α1T* or *Slo* in glioma flies reduced Ca^2+^ signaling to control levels (Fig. 4; Fig. S5). Furthermore, knocking down *Ca-α1T* or *Slo* did not significantly alter Ca^2+^ activity of non-tumoral glia (Fig. 4). Together, these findings show that Ca-α1T and Slo activities maintain elevated Ca^2+^ signaling in *Drosophila* glioma cells.

**Fig. 4.**
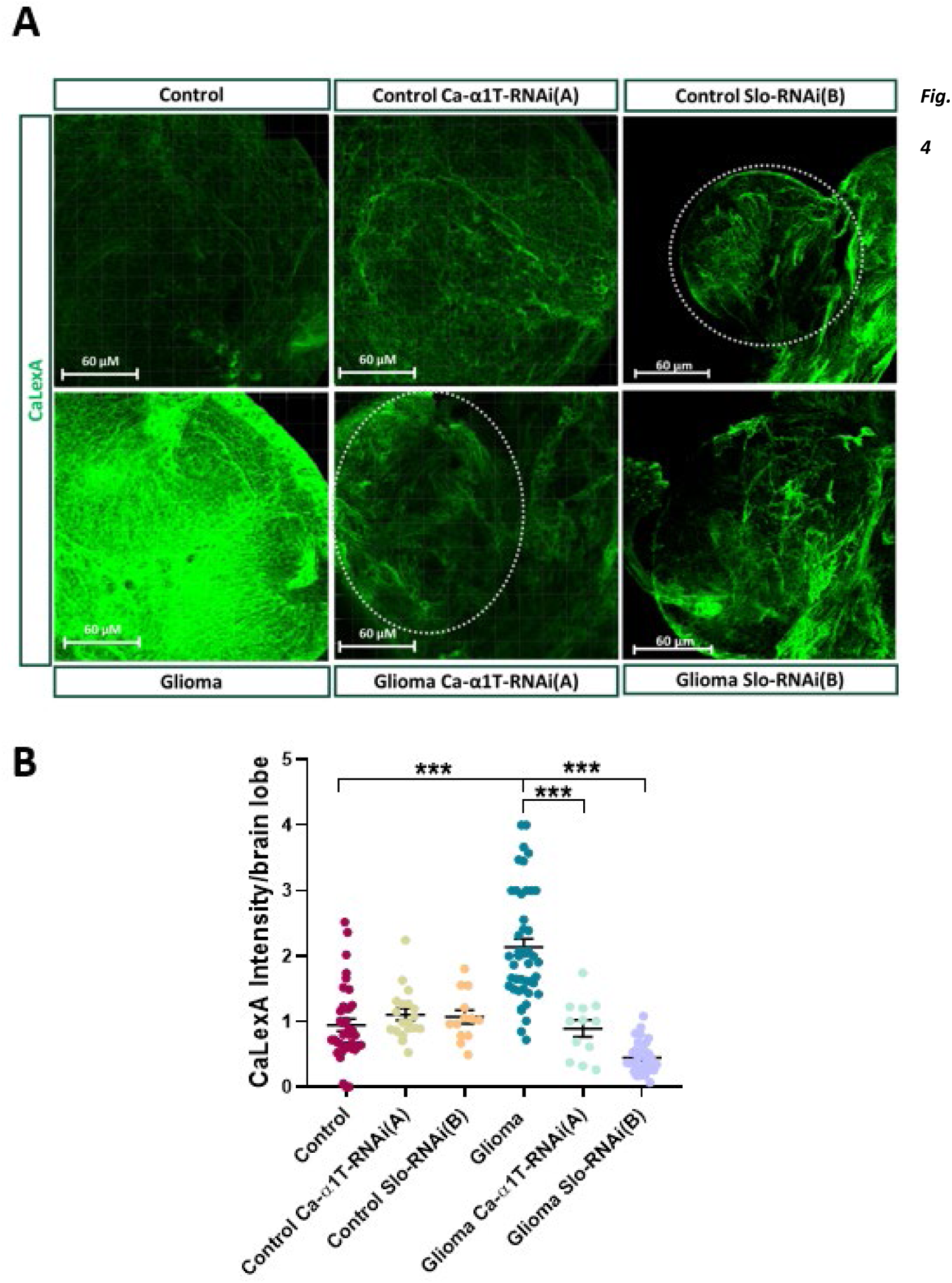
Knocking down Ca-α1T or Slo reduces the high cytoplasmic Ca^2+^ activity characteristic of gliomas. **(A)** Representative confocal images showing GFP fluorescence indicating glial Ca^2+^ activity via the GFP-CaLexA construct in Drosophila larval glioma or control brain lobes. Scale bar: 60 µm. **(B)** Quantification of GFP fluorescence intensity corresponding to glial Ca^2+^ activity. Gliomas exhibited significantly higher glial Ca^2+^ activity than control brains. Knockdown of Ca- α1T or Slo in glioma significantly reduces glial Ca^2+^ activity compared to control glioma. No significant changes were observed when Ca-α1T or Slo are knocked down in control brains. Values represent mean and SEM of GFP intensity per pixel and brain lobe. Data obtained from more than three independent experiments, with sample sizes between n=6-40 brain lobes (Ca-α1T) and n=8-37 brain lobes (Slo). One-way ANOVA with Tukey’s multiple comparisons test (***p<0.001).

### Reducing *Slo* expression, but not that of *Ca-a1T*, improves survival and relieves neurodegeneration of glioma flies

It has been previously shown that flies bearing gliomas have shorter life spans than control flies (Fig. 5A and D) [22]. As the expression of *Ca-α1T* or *Slo-*RNAi in glioma cells reduced cell proliferation and signaling, we evaluated whether it also impacted the survival of adult flies. *Ca-α1T-*RNAi expression in glioma cells did not significantly improve the survival of the adult flies, either males or females (Fig. 5A and C; Fig. S6). When these constructs were used to knockdown *Ca-α1T* in control flies, no differences in survival were observed, except for *Ca-α1T*- RNAi(A) that decreased survival of control flies (Fig. 5A and C).

**Fig. 5.**
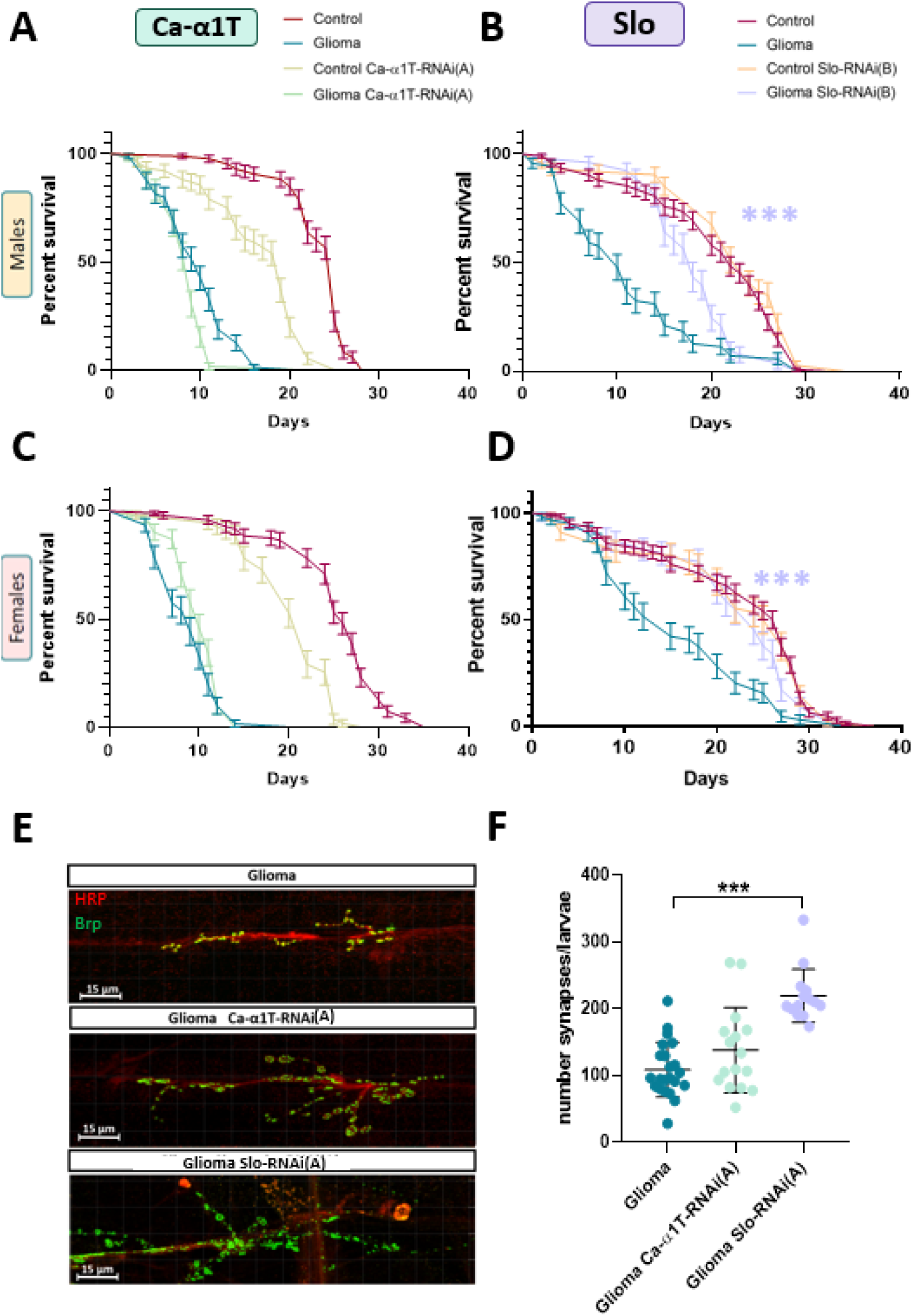
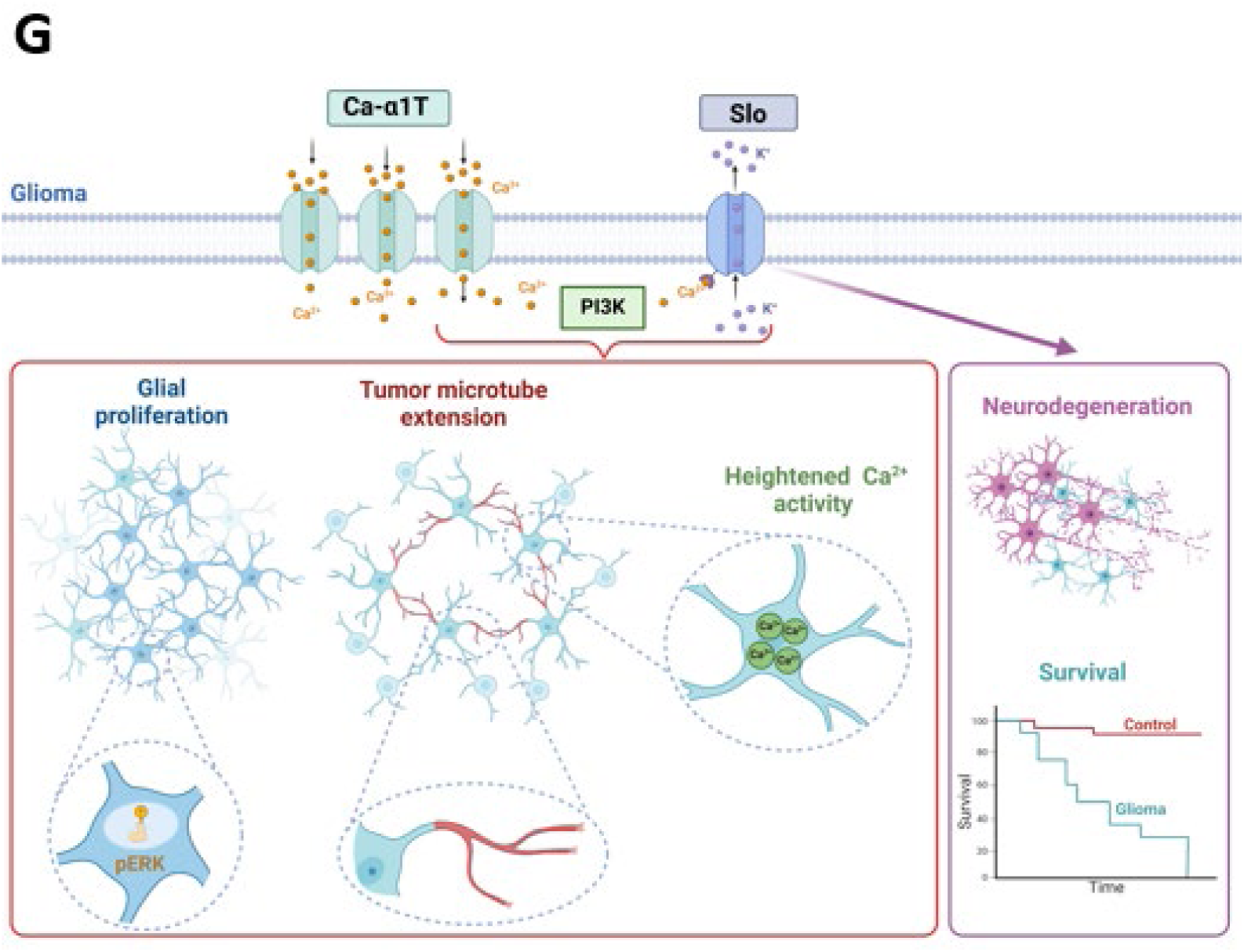
Knocking down Slo improves survival and increases NMJ synapses of glioma flies. Kaplan-Meier survival curves for adult male and female flies displaying the four experimental conditions. **(A-C)** Knockdown of Ca-α1T using Ca- α1T-RNAi(A) construct in glioma flies did not improve survival, while it decreased survival of control flies. **(B-D)** Knockdown of Slo using Slo-RNAi(A) significantly improves survival of glioma flies. In control flies, reducing Slo expression did not affect survival. Data from 8-9 independent experiments and a sample size between n=129-322 individuals per condition. Statistical analysis: Gehan-Breslow-Wilcoxon test. (***p<0.001). **(E)** Confocal microscopy images of synapses at the NMJ of Drosophila larvae with gliomas (neuronal membranes were stained in red with anti-HRP antibody and synaptic boutons were immunostained in green with anti-Brp). Scale bar: 15 μm. **(F)** Quantification of NMJ synapse number. Knocking down Ca-α1T in glioma did not significantly change synapse number compared to control glioma, whereas knocking down Slo in gliomas increased synapse number compared to control glioma larvae. Values represent means and SEM from more than three independent experiments (sample size between n=7-12 brain lobes). One-way ANOVA with Tukey’s multiple comparisons test (***p<0.001). **(G)** Proposed model: Ca-α1T and Slo channels regulate crucial aspects contributing to glioma growth like glial cell proliferation, TM extension, elevated Ca^2+^ activity and participate in ERK and PI3K signaling (features in the red box increase in the tumor and revert to control values upon channel knockdown). Adult glioma flies with reduced Slo levels show improved survival and signs of functional recovery (increased NMJ synapses; purple box).

However, reducing *Slo* expression extended the survival of glioma-bearing flies. The *Slo*- RNAi(A) construct significantly increased the survival for both males and females (Fig. 5B and D) and a second construct, *Slo*-RNAi(B), showed a modest improvement in the survival of glioma flies, particularly during the first 15-20 days (Fig. S6). Importantly, knocking down *Slo* in control flies did not alter survival with either RNAi construct (Fig. 5B and D; Fig. S6). Therefore, while targeting *Ca-α1T* and *Slo* reduced glioma cell proliferation, only *Slo* downregulation translated into a survival benefit of glioma-bearing individuals.

In order to address the difference in adult survival of flies bearing gliomas expressing *Ca-α1T* or *Slo-*RNAis, we decided to analyze the number of synapses at the neuromuscular junction (NMJ) as a measure to assess the level of glioma-induced neurodegeneration. We evaluated the number of NMJ synapses of segments A2, A3 and A4 in 3rd instar larvae by immunostaining the presynaptic neuronal membrane with anti-HRP antibodies [41] and the synaptic bouton using antibodies against the active zone protein Bruchpilot (Brp). *Ca-α1T-*RNAi expression did not alter NMJ numbers of glioma vs. control larvae (Fig. 5E and F). In contrast, *Slo-*RNAi expression in gliomas significantly increased the number of NMJs compared to the control glioma condition (Fig. 5E and F).

To determine if the increased synapse number at the NMJ translated into functional improvements, we analysed locomotion using a climbing assay. Consistent with the NMJ results, reducing *Slo* expression led to an improved locomotion of glioma-bearing flies, with a significant effect with *Slo*-RNAi(A) (Fig. S7). Locomotion of *Ca-α1T* RNAi-glioma bearing flies did not change compared to the control glioma conditions (Fig. S7). In conclusion, the correlation between increased NMJs number and enhanced locomotor ability suggests that reducing *Slo* expression mitigates glioma-induced neurodegeneration, thus improving the survival of glioma flies.

### Transcriptomic analysis reveals that knocking down *Ca-α1T* affects carbohydrate metabolism, whereas downregulating *Slo* reduces glutamatergic signaling in *Drosophila* glioma

Aiming at understanding the mechanisms behind the proliferative defects associated with *Ca- α1T* or *Slo* downregulation in gliomas and the improved survival of glioma flies with reduced *Slo* expression, we performed an RNAseq analysis on larval head samples. Four conditions (control, glioma, glioma *Ca-α1T*-RNAi(A) and glioma *Slo*-RNAi(A) were analyzed and differentially expressed genes were selected based on significant over- or under-expression in glioma compared to control, with a 2-fold change magnitude. From these, we identified genes whose expression returned to control levels upon knockdown, focusing on genes with known human orthologs (Fig. 6A). Next, to explore the biological processes behind the selected genes, we used the Gene Set Enrichment Analysis (GSEA) tool of Pangea and conducted Gene Ontology (GO) analysis (Fig. 6B-F). This approach allowed us to identify key cellular processes affected in glioma and rescued by *Ca-α1T* or *Slo* knockdown.

**Fig. 6.**
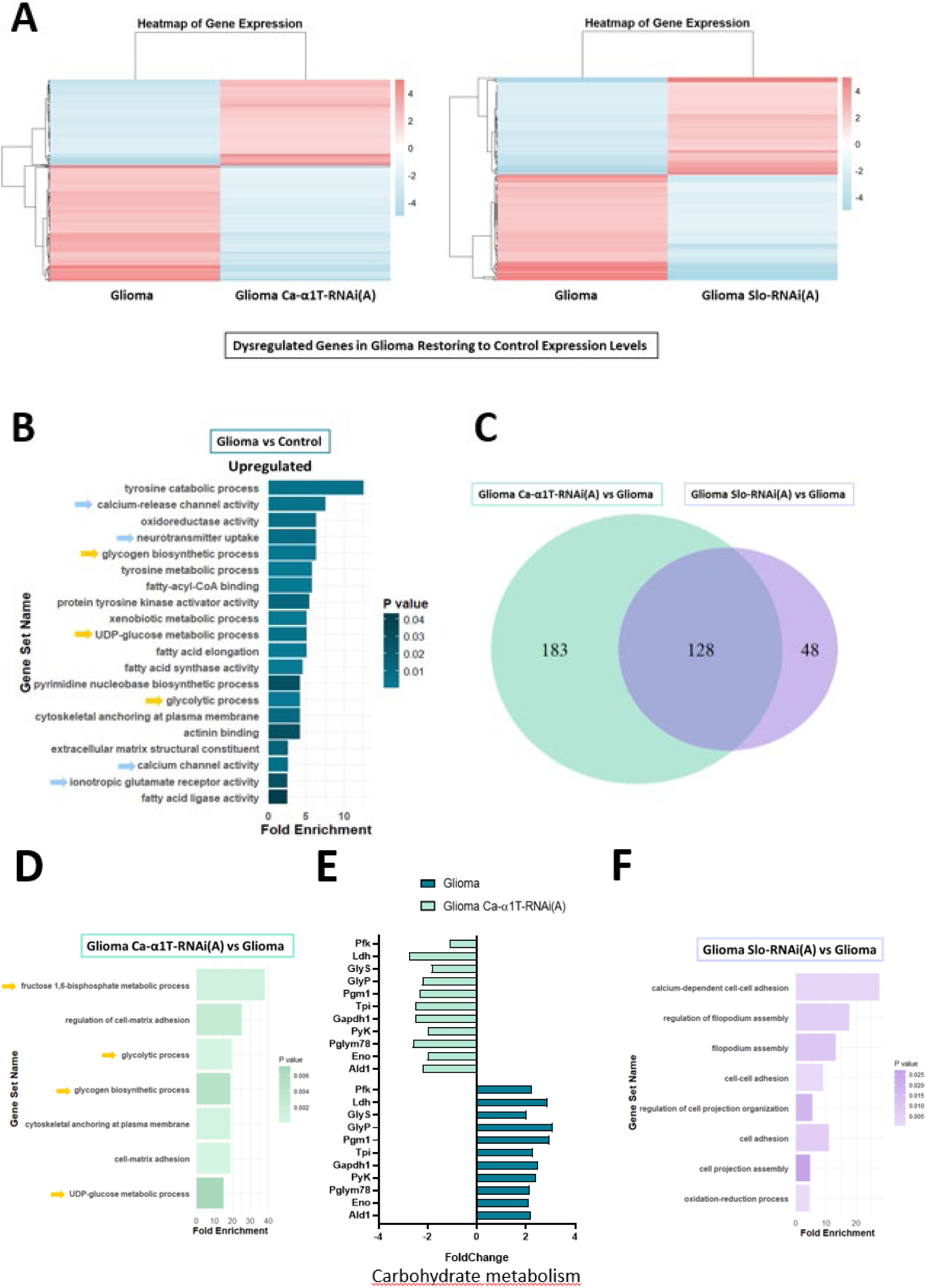
Transcriptomic analysis of gene expression changes and Gene Ontology categories in control gliomas and gliomas knockdown for Ca-α1T or Slo. **(A)** Heatmaps showing the expression of dysregulated genes in glioma that return to control-like levels upon knockdown of Ca-α1T or Slo. Enrichment Analysis (GSEA) of Gene Ontology (GO) categories related to biological processes using Pangea. (**B)** Biological processes overregulated in glioma vs control brain. Blue arrows highlight synaptic-related and neurotransmission-related activities, whereas yellow arrows label carbohydrate metabolic processes. **(C)** Venn diagram illustrates the number of protein-coding genes that are up- or down-regulated in glioma, returning to control-like values upon Ca-α1T or Slo knockdown. Genes specifically regulated by Ca-α1T-RNAi (183 genes in green) and Slo-RNAi (48 genes in purple) are highlighted, along with genes shared in the [43]knockdown of both channels (128 genes in the overlapping area). (**D and F)** Plots representing GSEA of GO categories performed using Pangea, based on the genes sets identified in the Venn diagram. Graphs show the most relevant GO categories in which these genes are involved, using the same colors as the Venn diagram. Fold Enrichment measures the proportion of GO term overrepresentation in the gene list compared to its occurrence in a reference set. P-values were calculated using Fisher’s exact test, with color gradient indicating statistical significance levels (p<0.05). **(E)** Genes associated to carbohydrate metabolism that reduced their expression to control values upon Ca-α1T targeting in glioma. RNAseq was performed on n=10 larval heads per condition. Results were adjusted using the Benjamini-Hochberg method (p<0.05).

Pathways upregulated in gliomas vs. controls were grouped into two classes, related to metabolism rewiring (oxidoreductase activity, amino acid, lipid or carbohydrate metabolism) or to synaptic activity (including neurotransmitter uptake, Ca^2+^ channel and ionotropic glutamate receptor activities) (Fig. 6B and Fig. S8). Additional GSEA analysis identified gene expression changes specifically affected by the knockdown of *Ca-α1T* or *Slo* in gliomas, or common to the knockdown of either channel (Fig. 6C). When looking specifically into gliomas expressing *Ca- α1T*-RNAi compared to control gliomas, categories related to cell-matrix adhesion and carbohydrate metabolism were downregulated (Fig. 6D). Genes linked to glycolysis and glycogen biosynthesis, whose expression significantly decreased in *Ca-α1T*-RNAi gliomas are shown in Fig. 6E. Targeting *Slo* in gliomas impacted on filopodium assembly and Ca^2+^ dependent cell-adhesion (Fig. 6F), with reduced levels of neuroligin 1 (*Nlgn1*) and *Ca-α1D*, a Ca^2+^ channel involved in glutamatergic synaptic plasticity (Fig. 7C).

**Fig. 7.**
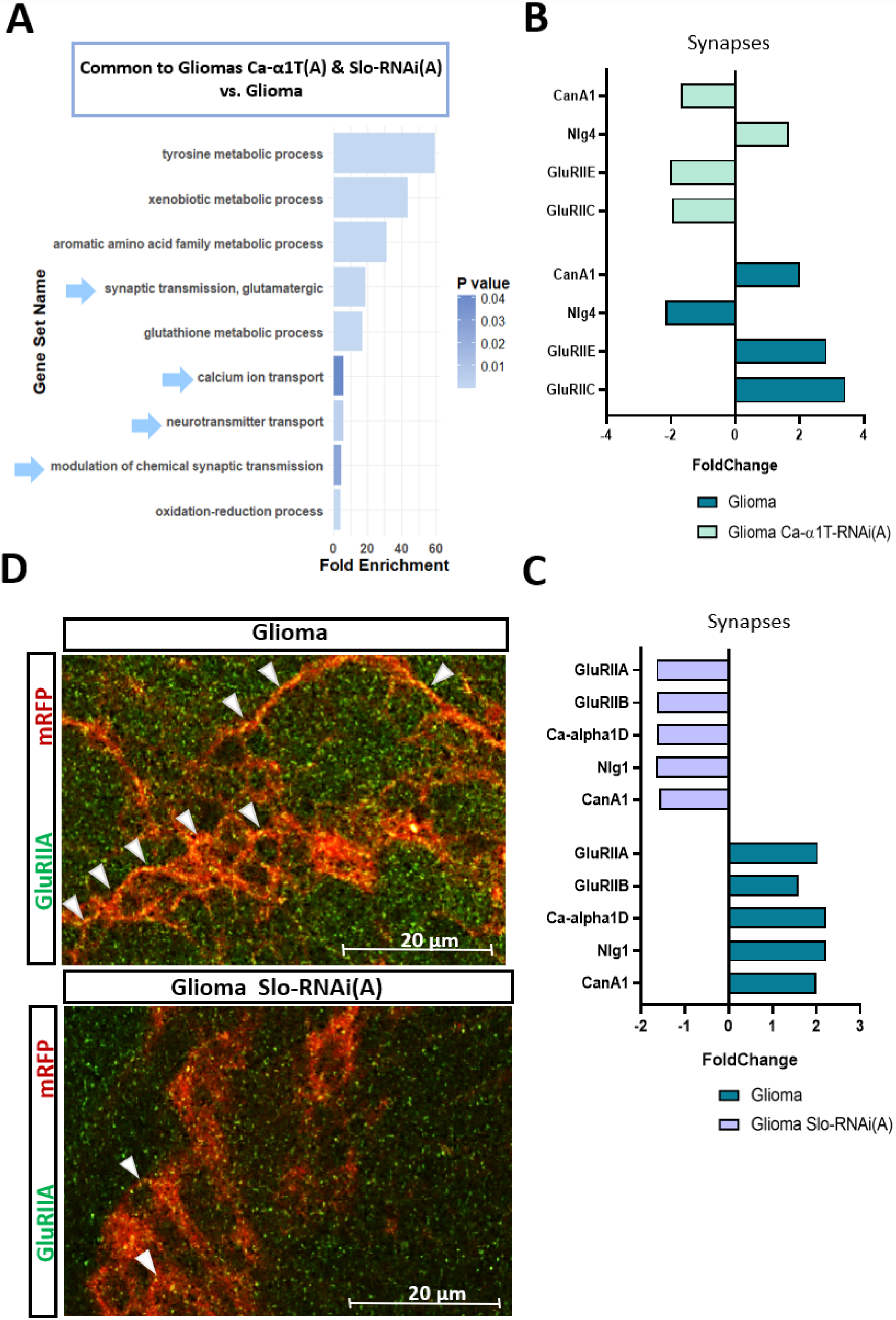
Expression of genes related to synapse function in Ca-α1T- or Slo-knocked down gliomas compared to control gliomas. **(A)** Graph shows the most relevant GO categories shared in Ca-α1T- or Slo-knocked down gliomas (blue arrows highlight synaptic-and neurotransmission-related activities). Fold Enrichment measures the proportion of GO term overrepresentation in the gene list compared to its occurrence in a reference set. P-values were calculated using Fisher’s exact test, with color gradient indicating statistical significance levels (p<0.05). (**B-C)** Genes associated to glutamatergic synapses that were dysregulated in glioma upon Slo or Ca-α1T knockdown, compared to control gliomas. Only genes that completely reverted their expression to control brain levels were selected. RNAseq results performed on n=10 larval heads/condition, were adjusted using the Benjamini-Hochberg method (p<0.05). **(D)** Immunostaining for GluRIIA decreases on glial mRFP-positive membranes of Slo-RNAi glioma cells compared to control glioma cells. Arrowheads point to yellow dots showing GluRIIA labelling on glial membranes. Bar= 20 μm.

Interestingly, common pathways downregulated in gliomas expressing RNAis for *Ca-α1T* or *Slo* vs. control gliomas included synaptic transmission and Ca^2+^ transport processes (Fig. 7A). Within the common downregulated synapse function category, *Ca-α1T* downregulation reduced *GluRIIC* and *GluRIIE* expression, that are structural subunits of AMPA receptors (Han et al., 2023) (Fig. 7B). However, knocking down *Slo* reduced the expression of AMPA glutamatergic receptor subunits *GluRIIA* and *GluRIIB* (Fig. 7C), both upregulated in gliomas and previously associated with tumoral growth. Decreased immunostaining of GluRIIA on glial membranes was confirmed in gliomas expressing *Slo*-RNAi compared to control gliomas (Fig. 7D). Thus, targeting *Ca-α1T* in *Drosophila* glioma affected carbohydrate metabolism, potentially contributing to reduced glioma cell proliferation. In contrast, targeting *Slo* in gliomas specifically reduced the expression of synaptic receptor subunits that may drive glioma overgrowth and aggressiveness, in agreement with the improved survival of the glioma flies with reduced *Slo* expression.

## Discussion

This work was designed to study the pro-tumoral roles of Ca-α1T (ortholog of mammalian CaV3) and Slo (ortholog of mammalian KCa1.1) in a *Drosophila melanogaster* glioma model. Both channels have independently demonstrated their relevance in GBM cell viability and invasiveness *in vitro* [8, 10, 17, 44]. However, most of the work targeting CaV3 and KCa channels in preclinical models has been performed using pharmacological blockers, for which numerous off-target effects have been reported [11, 25]. Here, we leveraged the extensive array of genetic tools and the high throughput -in terms of number of individuals- to study the role of these channels in a *Drosophila* glioma model. An additional advantage of this setting is the scarcity of channel isoforms, thus minimizing compensatory mechanisms that may arise following specific gene silencing.

### *Ca-*α*1T* and *Slo* regulate glioma cell proliferation, pro-tumoral pathways and elevate Ca^2+^ activity

Our data shows that *Ca-*α*1T* is overexpressed in the plasma membrane of glioma cells, recapitulating the observed upregulation of *CaV3.2* channel transcripts in human GBM biopsies (compared to normal brain tissue) and in glioma stem cells [44]. Moreover, in the 11-15% of patients presenting mutations that imply amplification of *CaV3.1* or *CaV3.2* channel genes, prognosis is impaired according to Kaplan-Meier survival estimates [14, 44]. Interestingly, a reciprocal positive regulation between effectors of the PI3K/Akt pathway and CaV3 channel expression/activity has been shown in different cell systems [11]. From this viewpoint, it is unsurprising that a glioma model based on the constitutive activation of PI3K is linked to *CaV3* channels overexpression. This argument stands for a significant proportion of human gliomas, as the *PI3KCA* gene (encoding the PI3K α subunit) presents gain of function mutations in 9% of IDH-wildtype and 12% IDH-mutant human gliomas [45]. In contrast, expression levels of *Slo* were similar in *Drosophila* glioma and normal glia cells, despite evidence for its heightened expression in a model of *dRAF* (ortholog of human BRAF)-gain of function *Drosophila* paediatric low-grade glioma model [19].

Targeting *Ca-*α*1T* or *Slo* with RNAi *in vivo* in *Drosophila* glioma validates relevant *in vitro* findings and reveals novel functions to be considered in a putative targeted therapy. Common to either channel knockdown we find: (1) a reduction of Repo+ glial cells in glioma brains, but not in brains of normal flies, indicating that these channels are mainly involved in the proliferation of transformed glia; (2) a reduction in glial membrane volume in glioma brains, which returns to normal brain levels. This measurement has been previously related to the formation of TM and part of the vampirization mechanisms of glioma cells on neurons [22]; (3) a reduction of glioma Ca^2+^ activity to the level of control brains, demonstrating a role for both channels in pro-tumoral Ca^2+^-dependent signaling; (4) reduced immunostaining for active pRII/p-ERK, which is inherently upregulated in our *Drosophila* glioma model downstream of EGFR constitutive activation [33]. Our data confirms the existence of a CaV3-ERK signaling axis in glioma cells previously reported in different cell settings [46–48]. It also confirms previous data regarding the association of KCa1.1 channels and ERK signaling, including reports of positive [19, 49] and negative [50] interactions; (5) a reduced PI3K activation (measured in control brains, since glioma brains express constitutively active PI3K). A positive relationship between PI3K activity and *CaV3.*1 expression has been shown in human GBM [10], prostate [51] and *PTEN*-deficient melanoma cells. Evidence linking KCa1.1 channels and PI3K activation was hitherto only pharmacological [52–54]. These findings are integrated in Fig. 5G.

### Knocking down *Ca-α1T* reduces the metabolic fitness of glioma cells, whereas targeting *Slo* decreases *GluRIIA and GluRIIB* expression and extends survival

RNAseq from larval heads was performed to gather further information about the pathways modulated by the ion channels object of our study. The expression of genes regarded as cuticular, pharyngeal or gustative markers was filtered out. We focused our gene ontology analysis on genes whose expression increases in glioma vs control flies, and that additionally return to control levels upon modulating *Ca-*α*1T o*r *Slo* channel expression.

This study identifies three main groups of processes activated in gliomas: (1) Metabolic rewiring: increased activities in amino acid, lipid and carbohydrate metabolism that support the high demands of proliferative gliomas [55]; (2) Cell adhesion, including Ca^2+^-dependent cell-cell adhesion and extension of cell protrusions or TM; (3) Elevated neurotransmitter receptor activity and Ca^2+^ signaling, potentially contributing to glioma growth and survival in a neural microenvironment. Shared and distinct effects were identified upon *Ca-*α*1T* or *Slo* knockdown across individual pathways.

Among other metabolic changes, knocking *Ca-α1T* specifically led to a reduction in glucose and glycogen metabolism, processes upregulated by the tumor [56, 57]. Moreover, *Ca-α1T*-RNAi reduces *Ldh* expression, responsible for the Warburg effect. Previous research has shown that ablating *LDH* isoforms in GBM mouse models reduces tumor growth, sensitizes tumors to radiotherapy and enhances survival. Similarly, the antiepileptic drug stiripentol, a LDH inhibitor, reduces GBM growth in mice [58]. Lactate accumulation has been recently linked with remodeling of the anaphase-promoting complex, thus directly impinging on the cell cycle machinery regulation [59]. Therefore, it is conceivable that decreased glial numbers in *Ca-α1T* RNAi-glioma flies are due to both decreased glycolysis and lactate production that fuel glial cell proliferation.

In addition, reducing *Slo* expression significantly downregulates proteins involved in filopodia assembly and Ca^2+^ dependent cell-cell adhesion, while targeting *Ca-α1T* downregulates cell-matrix adhesion proteins. Glioma cells form an electrically coupled network, synchronizing periodic Ca^2+^ oscillations that drive tumor progression [60]. Together, these downregulated GO categories by RNAi expression of either channel can explain the reduced glial membrane volumes in the RNAi-glioma fly strains, pointing to glioma cell communication defects upon channel modulation that reduce tumor growth.

Adult glioma-bearing flies with reduced *Slo* expression showed improved survival and signs of functional recovery (Fig. 5G). Thus, the reduction in glioma cell proliferation and glial membrane volume is not sufficient to alleviate neurodegeneration and improve survival of glioma-bearing flies, as the latter are observed only upon *Slo*-RNAi expression. Previous studies have shown that GBM cells participate in neuro-glioma synapses and constitute the postsynaptic element [32, 61]. Consistent with this, glioma cells express different GluRII subunits. Postsynaptic GluRs at the fly NMJs form heterotetramers comprised of the common core structural subunits GluRIIC, GluRIID and GluRIIE plus either the GluRIIA or the GluRIIB “signaling” subunits [43][62][63]. Our transcriptomic analyses show that *GluRIIA*, *GluRIIB*, *GluRIIC* and *GluRIIE* are upregulated in gliomas. *GluRIIA* and *GluRIIB* return to control levels in *Slo*-RNAi glioma flies, while *GluRIIC* and *GluRIIE* levels are normalized in *Ca-α1T*-RNAi flies. Interestingly, *GluRIIA* downregulation can reduce cytoneme formation [64] and improve survival of glioma flies [32]. These findings suggest that *Slo*-RNAi expressing glioma flies may increase their survival and rescue synaptic neurodegeneration caused by the tumor by reducing *GluRIIA* expression. *GluRII* orthologs in human are glutamate ionotropic AMPARs, which are being targeted in ongoing trials to interrupt the neuron-tumor connectivity and improve GBM prognosis [61][65][66]. The reduction of *Nlg1* (human *NLGN3 ortholog*) by *Slo* but not *Ca-*α*1T* channel knockdown in gliomas, also hints at a critical relationship between neurodegeneration and glioma fly lifespan. Activity-dependent secretion of synaptic Nlg3 is inversely associated with patient survival [67].

## Conclusions

Our work shows that both *Ca-*α*1T* and *Slo* channels are determinant for pro-tumoral, Ca^2+^- dependent signaling in *Drosophila* gliomas. The overlapping effects caused by their respective RNAi support a concerted action of both channels at the plasma membrane, as previously suggested [15], yet their distinct impacts on the survival of glioma-bearing individuals and some genomic features also reveal non-redundant roles. Overall, our data links increased fly survival and protection from neuronal degeneration to the downregulation of synaptic components whose expression is aberrantly elevated in glioma flies, highlighting this as a key therapeutic strategy. Thus, Slo channels prominently emerge as prospective therapeutic targets against GBM, warranting further research into their human orthologues, with particular attention to their widespread distribution in glia and neurons.

## Methods

### Fly stocks and genetic tools

Flies were bred in conventional fly food at 25°C. Any change in these conditions is specifically indicated. Fly stocks used were *UAS-lacZ* (BL8529), Oregon-R-C (BL0005), *repo-Gal4* (BL7415), *tub-gal80ts* (BL7019), *UAS-myr-RFP* (BL7119), *UAS-mLex-A-VP16-NFAT lexAop-rCD2-GFP* (CaLexA, BL66542), *UAS-Caα1T-RNAi* (BL39029), *UAS-Slowpoke-RNAi* (BL55405), *Tub>PH-GFP (BL8164) Mi{MIC}Caα1TMI08565* (BL44990) and *Mi{MIC}SloMI13492* (BL59344) from the Bloomington Stock Center (http://flystocks.bio.indiana.edu); *UAS-Caα1T-RNAi* (GD48008), *UAS- Cacophony-RNAi* (KK104178) and *UAS-Slowpoke-RNAi* (KK104421) from the Vienna *Drosophila* Resource Centre (https://stockcenter.vdrc.at); *UAS-dEGFRλ:UAS-dp110CAAX* were a gift of Dr. R. Read.

The Minos Mediated Integration Cassette (MiMIC) system was used to analyze the expression of the Ca-α1T and Slo channels in *Drosophila*. This system relies on a transposon containing a green fluorescent protein (GFP) cassette inserted downstream of the target gene promoter through site-specific integration [35]. The GFP signal allows visualization and quantification of Ca-α1T and Slo fusion protein expression levels in fixed tissues.

We utilized the CaLexA system, an NFAT-based calcium-dependent transcriptional reporter, under repo-Gal4 control to detect intracellular calcium activity specifically in glial cells. This system employs the LexA/LexAop gene expression system to drive GFP expression as a proxy for calcium activity [36].

We used a *Drosophila* line expressing the PH-GFP fusion protein under the ubiquitous tubulin promoter to monitor PI3K pathway activation. The PH domain specifically binds to PIP3, a lipid generated upon PI3K activation, leading to the recruitment of GFP to the plasma membrane. This translocation results in increased green fluorescence, used to assess PI3K activity [37].

### Drosophila glioma model

In *Drosophila*, coactivating dEGFR and dp110 (the catalytic subunit of PI3K) in glial cells gives rise to highly proliferative and invasive neoplastic cells, that mimic human gliomas and cause neurodegeneration [33]. To generate gliomas in *Drosophila* we used the Gal4/UAS system [38], with *Gal4* expression under the expression of the specific glial enhancer Repo and the glioma- forming mutations under UAS sequence (*repo-Gal4>UAS-EGFR^λ^, UAS-dp110^CAAX^*) [39]. Glioma- forming mutations were temporally regulated using the Gal80^TS^ repression system. When individuals were shifted to 29°C, Gal80TS underwent a conformational change, rendering it unable to bind to Gal4, thereby allowing their expression. Apart from the survival and locomotion experiments, in which flies were raised at 17°C and shifted to 29°C during adulthood, in the remaining experiments, the expression system was active during both embryonic and larval stages.

This system also allowed for the expression of a myristoylated form of the *red fluorescent protein* (*mRFP*) under the UAS sequence in glial cells of both glioma and control flies. Myristoylation facilitates the anchoring of mRFP to glial membranes, enabling the distinction between glial cells and neurons, and allows the quantification of glial membrane volume as an indirect measure of TM extension.

In this model, mutations triggering glioma are transcribed in glial cells simultaneously with the interfering RNAs (RNAi) targeting the ion channel of interest. Tub-Gal80TS; Repo-Gal4 and UAS- dEGFRλ:UAS-dp110CAAX flies were crossed with wild type flies (Oregon-R-C, BL0005) to generate control gliomas, with Slowpoke RNAi lines (*UAS-Slowpoke-RNAi(A)*, KK104421 and *UAS-Slowpoke-RNAi(B)*, KK104421) to generate gliomas with *Slowpoke* knockdown, and with Ca-*α*1T RNAi lines (*UAS-Caα1T-RNAi(A)*, GD48008 and *UAS-Caα1T-RNAi(B)*, BL39029) to generate gliomas with *Ca-α1T* knockdown.

### Immunohistochemistry

Brains of third-instar larvae were analyzed by immunohistochemistry. Brains were dissected, fixed in 4% formaldehyde + 0,3% Triton X-100 during 25 min and washed three times in phosphate-buffered saline + 0,3% Triton X-100 (PBT). Brains were blocked for 1h in BSA 5% PBT solution. Primary antibody was diluted in the blocking solution and incubated overnight at 4 °C. Then, primary antibody was washed thrice and secondary antibody in blocking solution incubated during 90 min at room temperature. Secondary antibody was washed with PBT and stained samples were mounted using Vectashield with DAPI (Vector Laboratories). Quantification of the number of glial cells was performed by counting Repo-positive cells per brain lobe.

For synapse quantification experiments, larvae were dissected, opened and fixed for 10 min in the same formaldehyde solution. The following steps were done as with brains. In these experiments, the synapses present in the neuromuscular junctions (NMJ) of segments A2, A3 and A4 of each larva were analyzed. These NMJs exhibit well-established morphological and physiological properties and are used to study glioma-derived neurodegeneration in

*Drosophila,* as the axons innervating these muscle pairs originate directly from the brain [21]. Primary antibodies used were: mouse anti-Repo (DSHB AB_528448, 1:300), mouse anti-GluRIIA (DSHB 8B4D2, 1:50), rabbit anti-GFP (Invitrogen A11122, 1:1000), goat anti-GFP (Invitrogen A11122, 1:1000), mouse anti-pRII/p-ERK (Merck Sigma-Aldrich M9692, 1:100), mouse anti- Bruchpilot (nc82, DHSB AB_2314866, 1:50), rabbit anti-HRP (Jackson ImmunoResearch 111- 035-144,1:400).

Secondary antibodies were: anti-mouse AlexaFluor 488 (Invitrogen A11001, 1:300), anti-rabbit AlexaFluor 594 (Invitrogen A21207, 1:300), anti-rabbit AlexaFluor 488 (Invitrogen A11008, 1:300) and anti-goat AlexaFluor 488 (Invitrogen A11055, 1:300). Antibodies against Slo were a kind gift of Dr. Xiaohang Yang (Zhejiang University, Hangzhou 310058, China).

### Imaging and staining quantification

Confocal images were acquired as serial optical sections every 1,51 μm using two different confocal microscopy systems: Leica Confocal Microscope TCS SP5 and Olympus Fluoview^TM^ FV1000. Brain lobe images for cell number and glial membrane analysis were obtained using 20x objective, while images of CaLexA analyses, synapse quantification, pRII immunostaining and PH-GFP were obtained using 40x objective with immersion oil. In the case of synapse quantification experiments, a digital zoom of 2x was added. MiMIC images were acquired using a 60x objective with immersion oil.

Brain lobe images are presented as Z-project images (all slices), except for pRII, PI3K, GluRIIA, Slo and MiMC in which a representative slice was chosen and magnified to facilitate visualization.

Images were analyzed using IMARIS® (Imaris 6.3.1 software) or Fiji (ImageJ 1.52v) depending on the experiment. Number of Repo-positive cells and mature active zones (nc82-stained synapses) were analyzed using IMARIS® spot counter tool. We established a minimum size and threshold for the puncta in the control samples of each experiment. Subsequently, these criteria were applied to analyze each experimental sample. Glial network volume was quantified using IMARIS® surface tool. MiMIC experiments were quantified using an ImageJ macro that analyzes pixel intensity in green over red region, equivalent to *Ca-α1T* or *Slo* expression in glia (Supp. Table 1). CaLexA and PH-GFP experiments were quantified using an ImageJ macro that analyses mean green pixel intensity (equivalent to CaLexA or PH-GFP) (Supp. Table 2). Manual counting was employed for pRII experiments. Results were analyzed separately for each brain lobe, except for the synaptic quantification experiments, which reported the mean number of synapses per larva.

### Survival assay

Parental lines carrying *Tub-Gal80TS; Repo-Gal4* and *UAS-dEGFR^λ^:UAS-dp110^CAAX^*were respectively crossed with *UAS-LacZ* (control) or *UAS-RNAi*. Progeny individuals obtained formed the conditions control, glioma, control+RNAi and glioma+RNAi and were put at 29°C and survival was calculated as the percentage of surviving flies respect to the starting number. Each experiment was performed at least 4 times, using vials that carried 10 flies separated into male and female groups. Alive individuals were counted each 2-3 days.

### Climbing assay

Locomotor ability in flies was assessed using the climbing assay, which leverages their natural behaviour of negative geotaxis (tendency to move upward against gravity). The same genetic crosses and individuals used in survival assays were tested in the climbing assay.

Flies were placed in an empty and sealed tube marked with a 5 cm line from the bottom. Flies were tapped to the bottom of the tube and the number of flies climbing above the mark within five seconds was recorded. Assay was performed every 2 to 3 days, which each session comprising 8 replicates conducted on the same day. The entire experiment was repeated at least four times to ensure reproducibility.

### Real time PCR

To determine the efficiency of RNAi-induced silencing, a *Drosophila* line ubiquitously expressing *Gal4*, *Gal80^TS^*, and *GFP* under the control of the tubulin promoter and UAS sequence was used. To ensure accurate quantification of silencing, larvae with ubiquitous RNAi expression were selected, avoiding potential contamination from glia-specific Gal4 models, where dissecting brains could include neurons not expressing RNAi. Flies carrying various RNAi constructs were crossed with this line, and progeny displaying RNAi-induced silencing across all tissues were chosen. RNA was extracted from pools of 5–10 whole larvae for quantitative PCR analysis.

RNA was isolated using the EZNA Total RNA Kit I (VWR, R6834-01). One microgram of RNA was reverse transcribed at 25°C for 10 min, 42°C for 1 h, and 92°C for 5 min using the RevertAid RT Reverse Transcription Kit (VWR, Radnor, PA, USA; K1622). The resulting cDNA was analyzed by quantitative PCR (qPCR) using a CFX96™ Real-Time PCR Detection System (Bio-Rad, Hercules, CA, USA) with TaqMan hydrolysis probes labelled with FAM and TaqMan™ Gene Expression Master Mix (Thermo Fisher Scientific, 10525395). Gene-specific probes included *Drosophila Ca- α1T* (Dm01796209_g1), *Slo* (Dm02150795_m1), and *RplII* (internal control, Dm01813383_m1), all from Thermo Fisher Scientific. Each gene was analyzed in triplicate, and relative expression was calculated using the ΔΔCt method (Applied Biosystems) and plotted.

### RNAseq

Total RNA was extracted from *Drosophila* samples using the EZNA Total RNA Kit I (VWR, R6834- 01), using DNase treatment to eliminate potential DNA contamination. RNA sequencing (RNAseq) was performed by Macrogen. For mRNA library preparation, the TruSeq Stranded mRNA Library Prep Kit was used, and sequencing was carried out on an Illumina platform, generating 151 bp paired end reads. Raw sequencing data underwent quality control using FastQC, and low-quality reads and adapter sequences were removed with Trimmomatic. The resulting reads were aligned to the *Drosophila* reference genome (dm6 version) using HISAT2, a splice-aware aligner. Transcript assembly and gene expression quantification were subsequently performed with StringTie. Expression levels were calculated as read counts, FPKM (Fragments Per Kilobase of transcript per Million mapped reads), and TPM (Transcripts Per Million).

Differentially expressed genes (DEGs) were identified using edgeR in 10 pairwise comparisons. Genes with an absolute fold change ≥ 2 and an adjusted p-value (pvalbh) below 0.05 were considered differentially expressed. The company provided an excel file with gene expression data, including normalized values (FPKM and TPM), read counts per gene, fold change, and adjusted p-values calculated using the Benjamini-Hochberg method to control the false discovery rate.

The following comparisons were analyzed: Glioma vs. Control, Glioma *Ca-α1T*-RNAi(A) vs. Glioma, Glioma *Slo*-RNAi(A) vs. Glioma, Glioma *Ca-α1T*-RNAi(A) vs. Control, and Glioma *Slo*- RNAi(A) vs. Control. Differential expression analysis identified genes with absolute fold changes >2 or <−2 and adjusted p-values (pvalbh) <0.05, considered significantly overexpressed or underexpressed. In the comparison Glioma *Ca-α1T*-RNAi(A) vs. Control and Glioma *Slo*-RNAi(A) vs. Control, genes with fold changes between −2 and +2 and non-significant pvalbh values were considered to have reverted to control-like expression levels upon silencing, thus highlighting genes potentially involved in restoring normal cellular states. Uncharacterized genes, non-protein-coding genes, and *Drosophila* genes lacking a known human ortholog were excluded to focus on functional and translationally relevant targets.

### Gene Set Enrichment Analysis (GSEA) and Gene Ontology (GO) Analyses

Gene set enrichment analysis was conducted using the PANGEA platform (PAthway, Network, and Gene-set Enrichment Analysis) with separate lists of significantly upregulated and downregulated genes. GO term enrichment was determined by comparing their frequencies to reference datasets, highlighting relevant biological processes. Statistical significance was assessed with Fisher’s exact test and adjusted using FDR or Bonferroni corrections. GO category lists were visualized using R Studio, excluding redundant, overly general, or *Drosophila*-specific terms to focus on relevant biological processes. This approach provided detailed insights into the molecular mechanisms underlying the observed phenotypes.

## Acknowlegments

This work was funded by the Spanish Ministry of Science and Innovation (Programa RETOS RTI2018-094739-B-I00 and Generación de Conocimiento PID2022-138688OB- I00 to CC and JH; PID2022-139786OB-I00 and PI22CIII/00062 to SCT, and PID2022-141323NB-I00 to AC) and by Fundació La Marató de TV3 (201909-30 to CC). We thank Alba Vicario, Laia Castells and Diana Alexandra Anitoaei for help maintaining fly stocks. We also acknowledge the support of Anaïs Panosa and the UdL Confocal Microscopy. LA was a recipient of an IRBLleida/Diputació de Lleida and FI-Agaur predoctoral fellowships. PM-L was a recipient of UdL predoctoral fellowship.

## Data availability

The RNAseq datasets generated during the current study are submitted and will be available in the Gene Expression Omnibus upon publication of the manuscript. The rest of the data will be available upon reasonable request.

## Author contributions

The first author and all senior authors contributed to the study conception and design. Material preparation, data collection and analysis were performed by Lía Alza, Patricia Montes-Labrador, Diego Megías, Sergio Casas-Tintó, Andreu Casali, Judit Herreros and Carles Cantí. The first draft of the manuscript was written by Judit Herreros and Carles Cantí and all authors commented on previous versions of the manuscript. All authors read and approved the final manuscript.

The authors have no relevant financial or non-financial interests to disclose.

## Supplementary Figures

**Supplementary Fig. 1.**
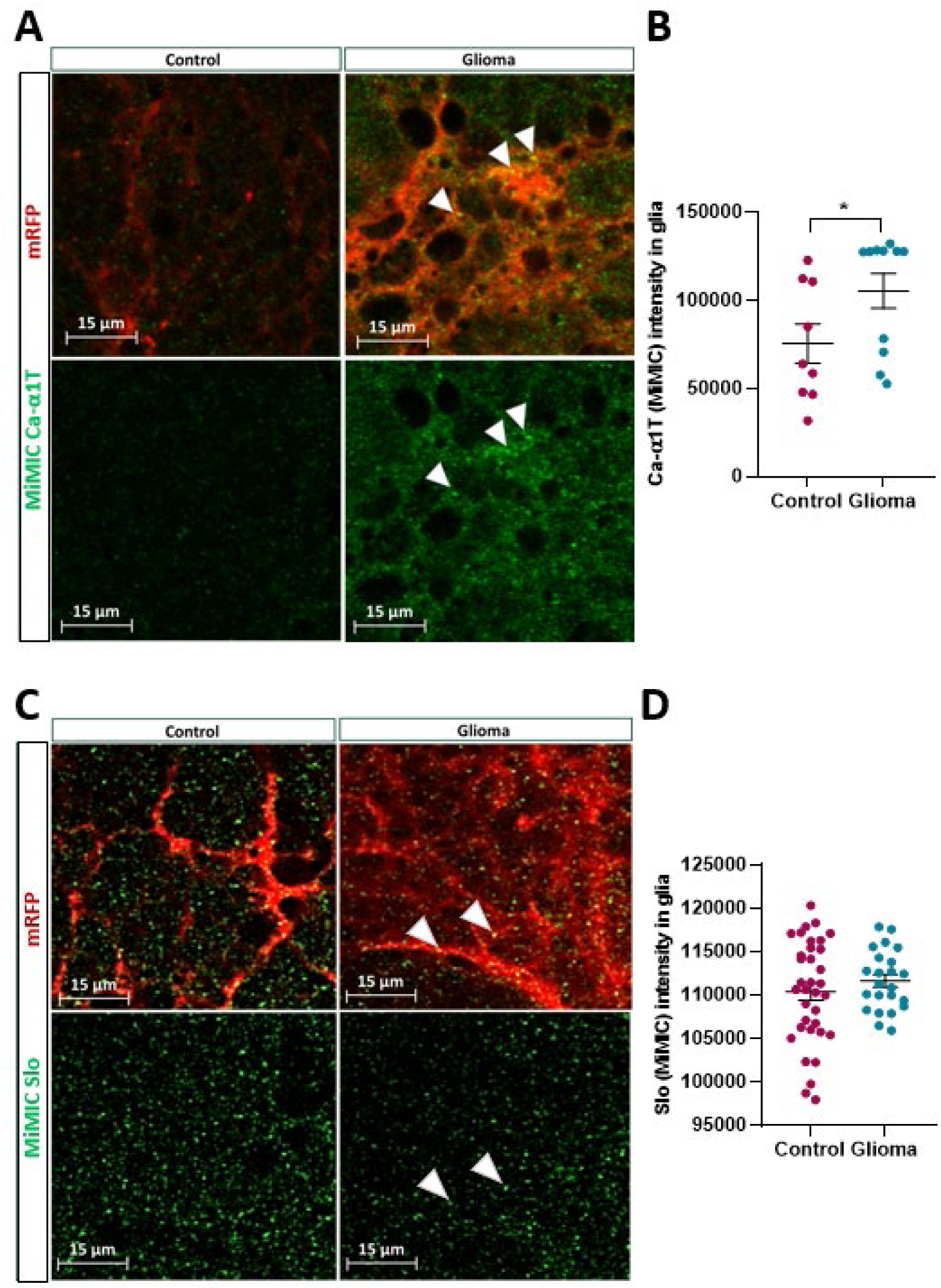
Expression analysis of Ca-α1T and Slo by using MiMIC tools in Drosophila glioma vs. control brain. **(A-C)** Representative confocal images showing **(A)** Ca-α1T or **(C)** Slo expression reported in green (GFP) using specific MiMIC tools. mRFP corresponds to glial membranes; green signal over red signal (yellow dots) indicates GFP-channel fusion protein on glial membranes. Scale bar: 15 µm. **(B)** Quantification of GFP showing Ca-α1T expression, which is significantly increased in tumoral glia compared to control glia. **(D)** Quantification of GFP showing Slo expression in tumoral vs. control glia did not show significant changes. Values represent means and standard error of the mean (SEM), obtained from more than three independent experiments, with a sample size between n=10-38 brain lobes. (*p<0.05, T-test).

**Supplementary Fig. 2.**
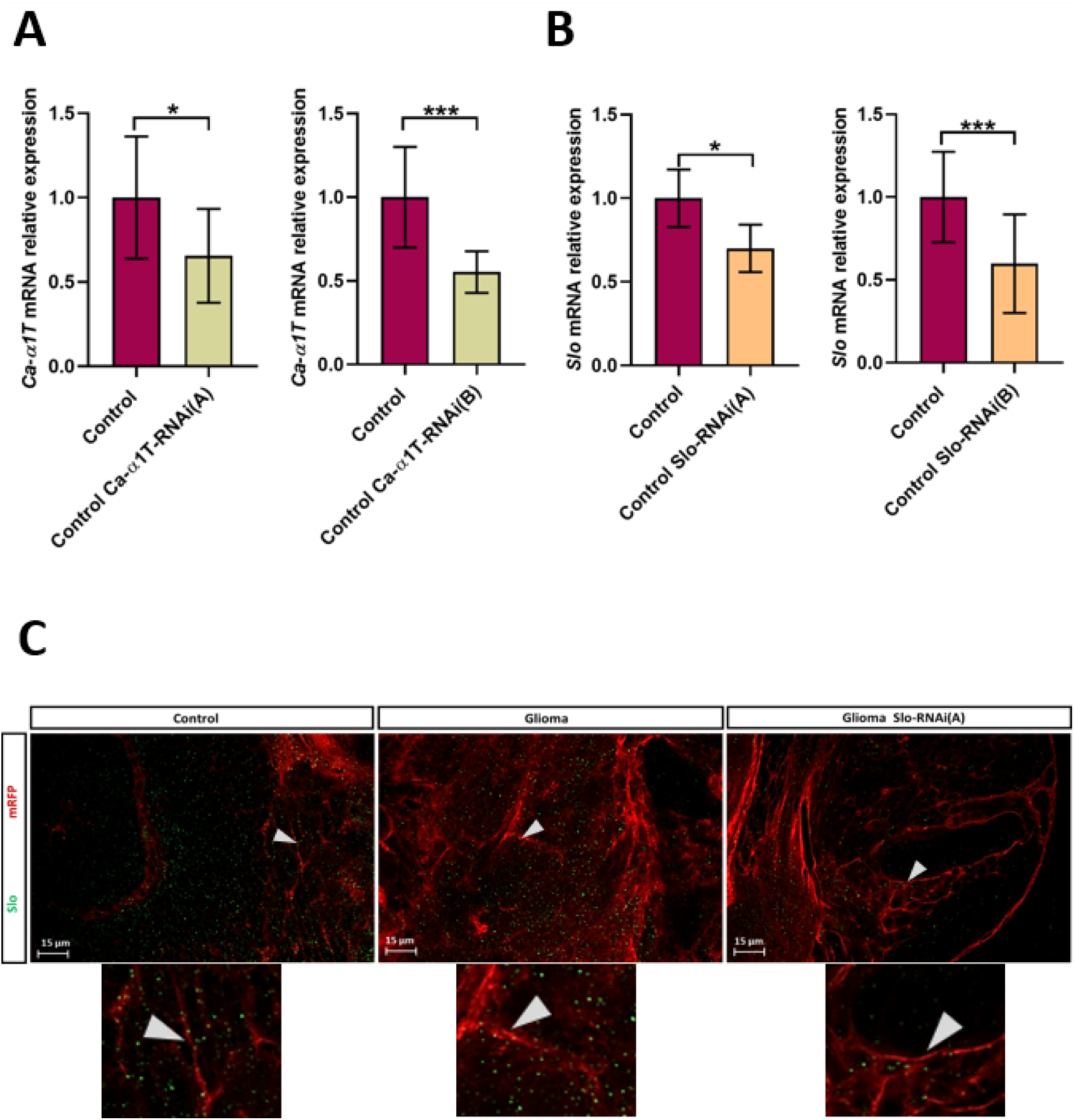
Effective gene knockdown by RNAi constructs from Drosophila lines targeting Ca-α1T or Slo. **(A-B)** qPCR analysis was performed in control larvae and larvae expressing RNAi sequences against Ca-α1T or Slo. RNAi constructs were expressed in all larval tissues under the control of tubulin promoter. Ca-α1T- RNAi(A) and RNAi(B) reduced the expression of Ca-α1T by 35% and 45%, respectively. Slo-RNAi(A) and RNAi(B) reduced the expression of Slo by 31% and 41%, respectively. Results are normalized to mRNA levels of RpL32. Values represent means and SEM. N=5-7 different experiments. T-test (***p<0.001, **p<0.01, *p<0.05). **(C)** Slo immunostaining (green) from control brain, control glioma and glioma expressing Slo-RNAi confirmed similar levels of Slo immunostaining between control and glioma brains and reduced immunostaining of Slo on mRFP- glial membranes in Slo-knockdown gliomas. Arrows point to Slo-positive puncta on glial mRFP membranes. Bar= 15 µm.

**Supplementary Fig. 3.**
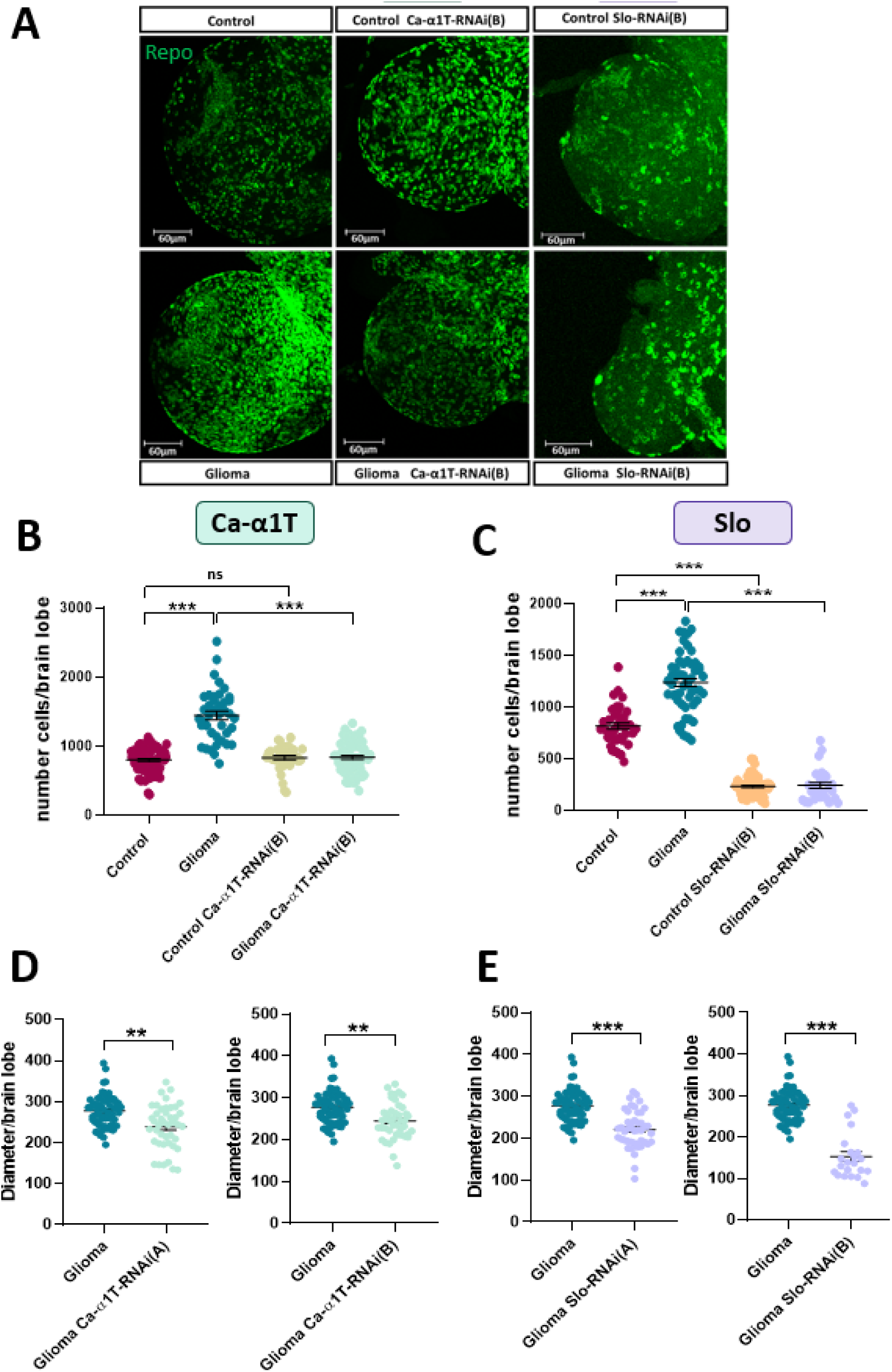
Ca-α1T or Slo knockdown reduces glioma cell proliferation in Drosophila. **(A)** Representative images of Repo-immunostained glial cells in Drosophila larval brain lobes. Scale bar: 60 µm. **(B- C)** Quantification of the number of glial cells per lobe, which increased in glioma vs. control. Ca-α1T-RNAi(B) or Slo-RNAi(B) in glioma significantly decreased the number of glial cells vs. glioma. Ca-α1T knockdown using Ca- α1T-RNAi(B) in control brains showed no differences in cell numbers compared to control, while Slo-RNAi(B) reduced glial cell number in controls. **(D-E)** Glioma brain lobe diameters, either control and knockdown for Ca- α1T or Slo. Values represent means and SEM from more than three independent experiments, with a sample size n=21-88 brain lobes. One-way ANOVA with Tukey’s multiple comparisons test (***p<0,001, **p<0,01).

**Supplementary Fig. 4.**
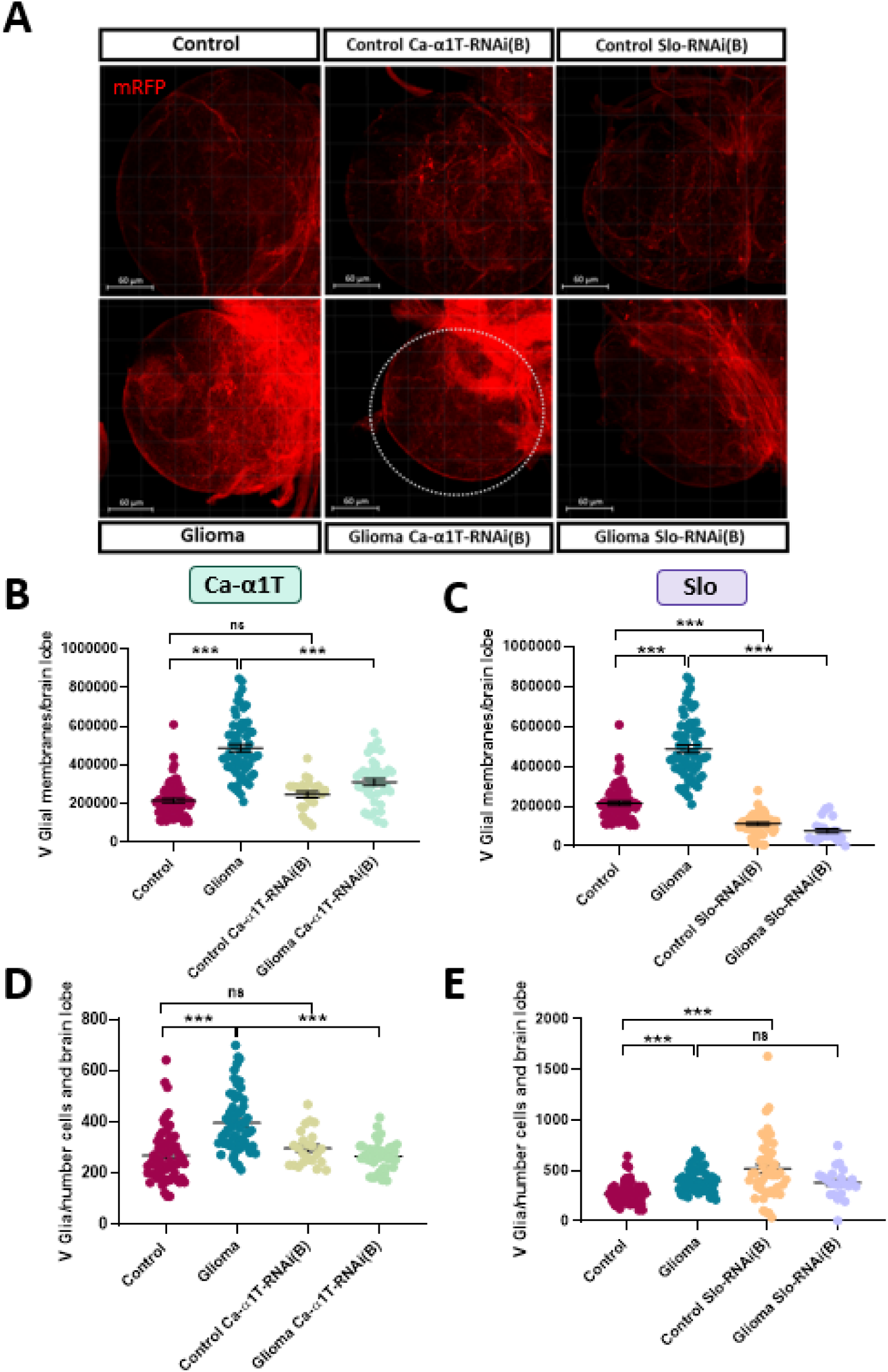
Ca-α1T or Slo knockdown reduces membrane extension in Drosophila glioma cells. (**A)** Representative confocal images of glial membranes stained for mRFP in Drosophila larval brain lobes. Scale bar: 60 µm. **(B-C)** Quantification of glial membrane volume per brain lobe. Gliomas showed significantly higher membrane volume compared to control. Ca-α1T-RNAi(B) and Slo-RNAi(B) significantly decreased membrane volume in gliomas. Knockdown of Ca-α1T in controls did not affect membrane volume, while Slo-RNAi(B) significantly decreased it. **(D)** Ca-α1T-RNAi(B) reduced membrane volume per cell in glioma. (**E)** Slo-RNAi(B) did not decrease membrane volume per cell in glioma, which is attributed to a bigger effect in cell proliferation by this construct (Supp. Figure 3C). Data represents means ± SEM from at least three independent experiments, with a sample size between n=11-76 brain lobes per condition. One-way ANOVA with Tukey’s multiple comparisons test (***p<0.001).

**Supplementary Fig. 5.**
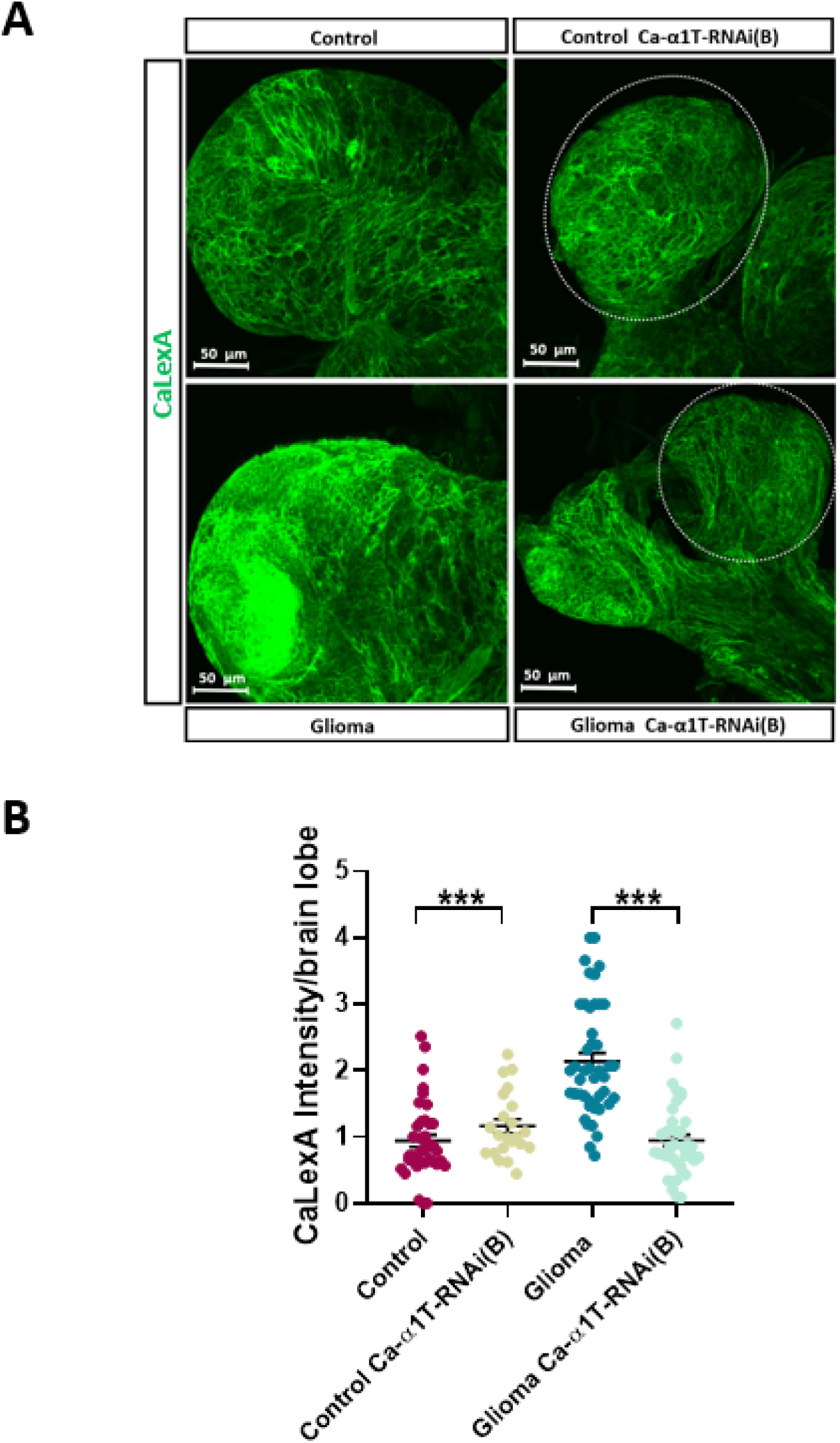
Ca-α1T knockdown using Ca-α1T-RNAi(B) construct reduces the elevated Ca^2+^ activity characteristic of glioma. **(A)** Representative confocal images showing GFP fluorescence indicating glial Ca^2+^ concentrations via the GFP-CaLexA construct in Drosophila larval brain lobes. Scale bar: 50 µm. (**B)** Quantification of GFP intensity corresponding to glial Ca^2+^ activity. Glioma exhibited significantly higher glial Ca^2+^ activity than control brains. Ca-α1T-RNAi(B) construct decreased glial Ca^2+^ activity in glioma, without significantly affecting that in control brains. Values represent means and SEM obtained from more than three independent experiments, with sample sizes between n=6-40 brain lobes. One-way ANOVA with Tukey’s multiple comparisons test (***p<0.001).

**Supplementary Fig. 6.**
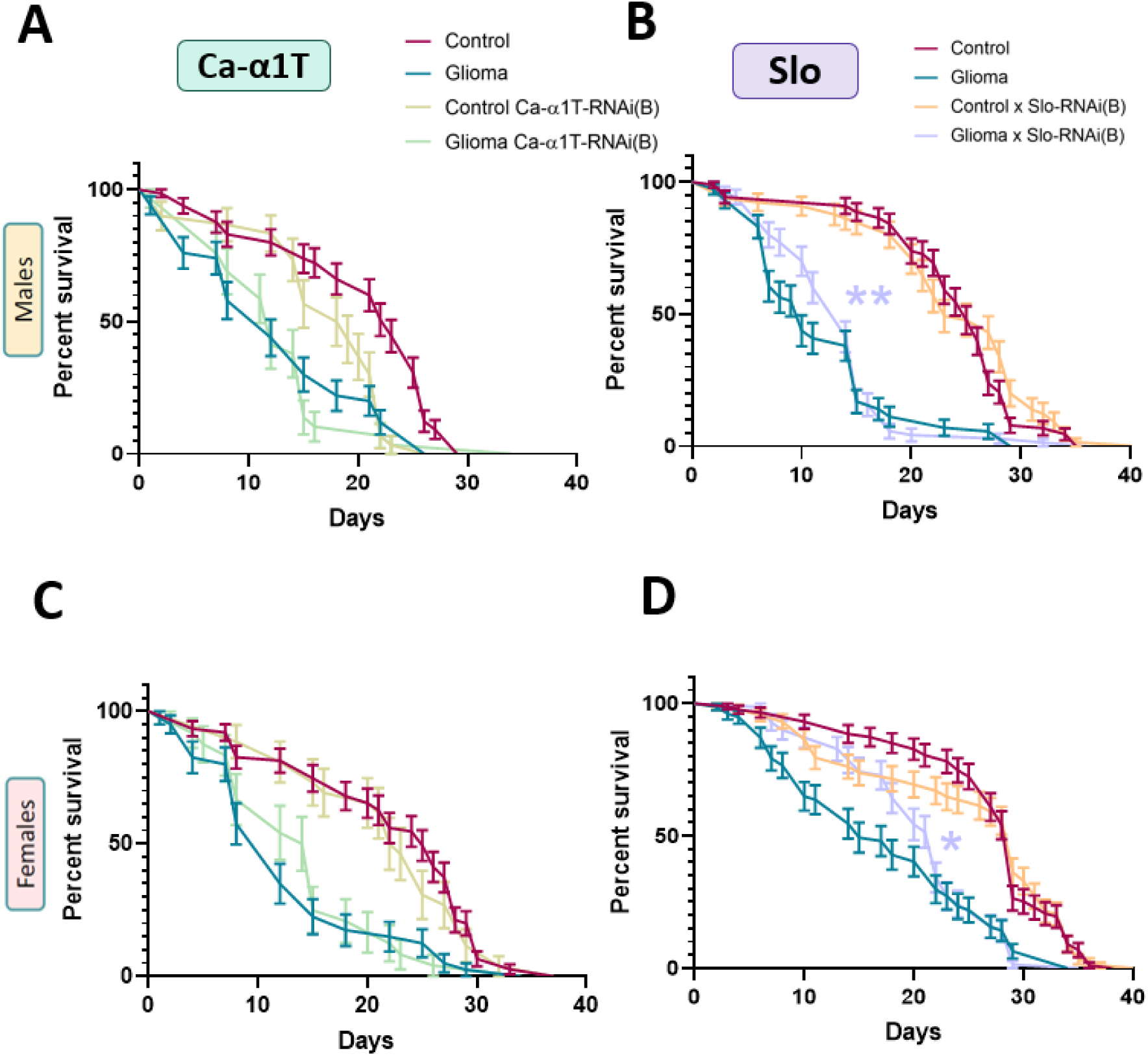
Slo knockdown improves survival of glioma-bearing flies. **(A-C)** Ca-α1T-RNAi(B) did not enhance survival in glioma-bearing flies and slightly affected survival of control males. **(B-D)** Slo-RNAi(B) improved survival of glioma-bearing flies. Gehan-Breslow-Wilcoxon test. test (**p<0.01; *p<0.05).

**Supplementary Fig. 7.**
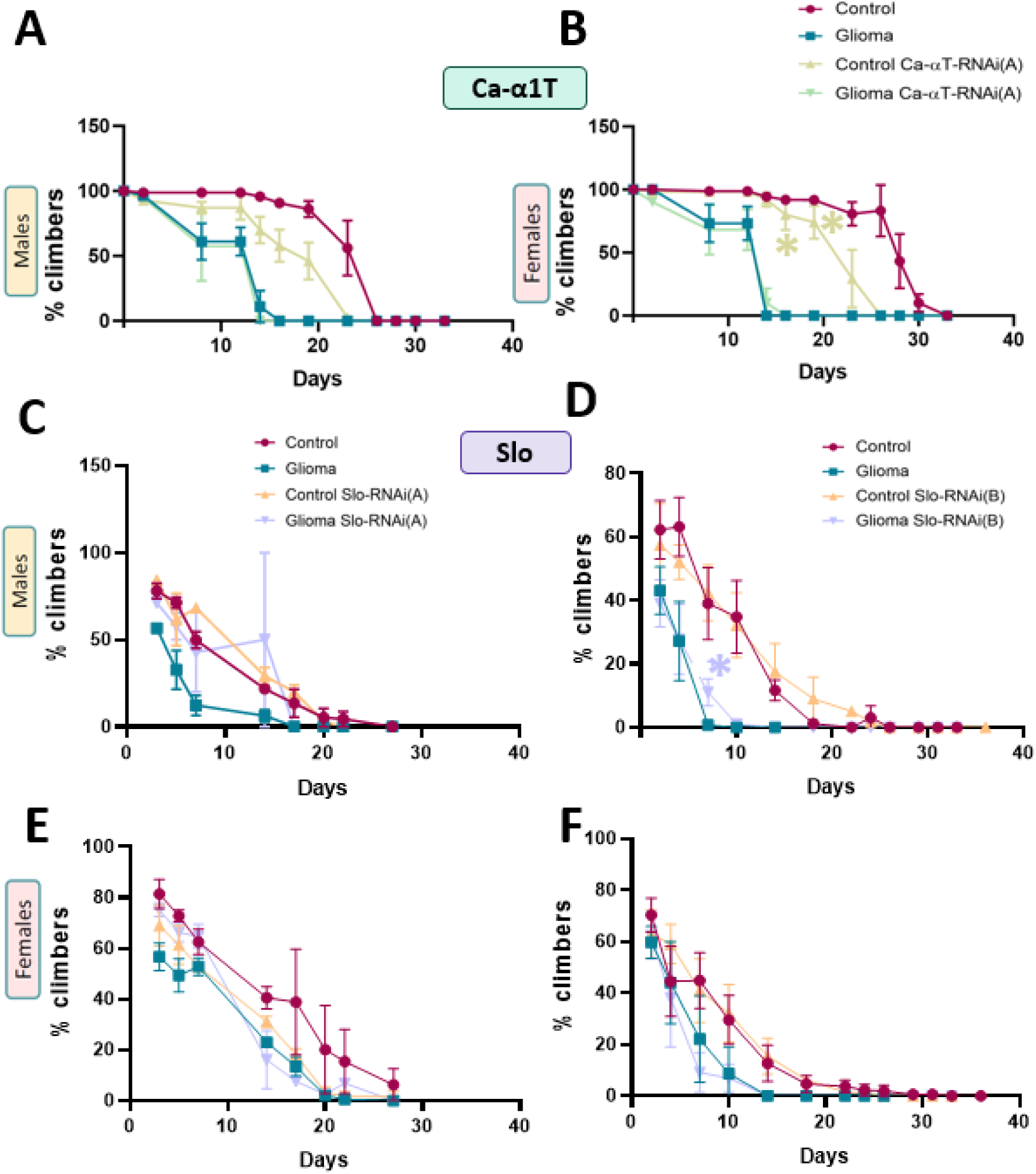
Knocking down Slo rescues locomotion deficits in male flies with glioma. Graphs show the percentage of flies climbing above a 5 cm line within 5 seconds. Glioma-bearing flies exhibited reduced climbing ability compared to controls. **(A-B)** Expressing Ca-α1T-RNAi(A) in glioma did not improve locomotion but significantly decreased locomotion of control females. (**C-D)** Knocking down Slo using Slo-RNAi(A) in glioma slightly improved locomotion in females during early days and significantly in males (*). (**E-F)** Slo knockdown using Slo-RNAi(B) in glioma slightly improved locomotion during early days and significantly in males. No differences were observed in controls. T-test performed by day (*p<0.05). Values represent means ± SEM from more than three independent experiments, total sample size n=68-322 individuals per condition.

**Supplementary Fig. 8.**
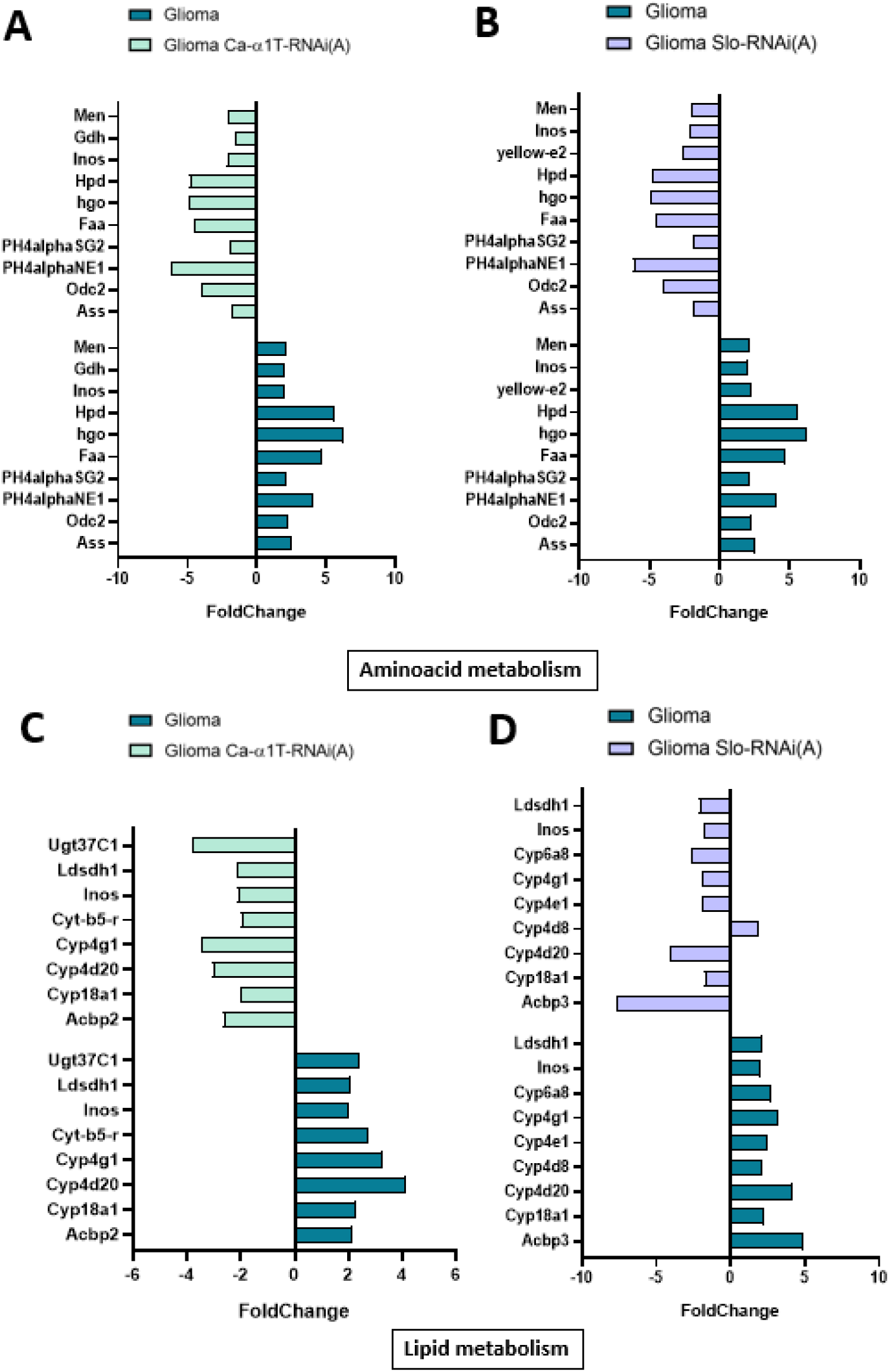
Expression changes in genes of the amino acid and lipid metabolism repressed following the Ca-α1T or Slo knockdown in gliomas. **(A-B)** Genes associated to amino acid metabolism that reduced their expression after Ca-α1T **(A)** or Slo **(B)** silencing. **(C-D)** Genes related to lipid metabolism that reduce their expression upon Ca-α1T **(C)** or Slo **(D)** knockdown. Only genes that completely reverted their expression to healthy brain levels were selected. RNAseq results, performed on n=10 larval head per condition, were adjusted using the Benjamini-Hochberg method (p<0,05).

**Supplementary Table 1.** ImageJ macro code for analyzing the mean pixel intensity of the green (GFP) channel over the red channel across each layer.

| Item | Description |
| --- | --- |
| Macro Name | **Masked_Fluorescence_Analysis.ijm** |
| Description | This macro analyzes the fluorescence intensity of a channel over a mask derived from another channel across multiple slices. |
| Author | Diego Megías |
| GitHub Repository | https://github.com/LiaAlzaBlanco/ImageJ-Macros-GFP-Analysis/blob/main/Masked_Fluorescence_Analysis.ijm |
| Requirements | ImageJ v1.54f, Bio-Formats Plugin v8.0.1, Java 1.8.0_322 |

**Supplementary Table 2.** ImageJ macro code for analyzing the mean pixel intensity of the green (GFP) channel across each layer.

| Item | Description |
| --- | --- |
| Macro Name | **Fluorescence_Intensity_Analysis.ijm** |
| Description | This macro analyzes the fluorescence intensity of a selected channel across multiple slices. |
| Author | Diego Megías |
| GitHub Repository | https://github.com/LiaAlzaBlanco/ImageJ-Macros-GFP-Analysis/blob/main/Fluorescence_Intensity_Analysis.ijm |
| Requirements | ImageJ v1.54f, Bio-Formats Plugin v8.0.1, Java 1.8.0_322 |

